# Generalization of the Packing Parameter for Quantifying the Morphology of Peptide Amphiphile Micelles

**DOI:** 10.1101/2024.06.10.598326

**Authors:** Luke E. Kruse, Bret D. Ulery, Karl D. Hammond

## Abstract

We present a quantitative means for classifying the shape of molecular dynamics simulated peptide amphiphile micelles (PAMs) that is both consistent with existing metrics and extendable to estimating shape-dependent free energy contributions. The presented framework not only outlines an approach for characterizing the shape of simulated PAMs but also presents expressions that can readily be applied to quantify the shape of particles from experimental techniques where aspect ratios are measured. The generalization of the packing parameter introduces a characteristic length that, when applied to simulated PAMs, functions intuitively as an effective radius for a PAM whose core is a perfect sphere or an infinite cylinder. The presented shape assignment scheme is used to develop a model for the free energy penalty associated with packing the tails of the amphiphiles into a core whose shape is modeled by an ellipsoid. Good agreement with previous models and scaling behaviors is observed and the importance of accounting for the shape and size dependence of the core is illustrated.

**TOC Graphic:** 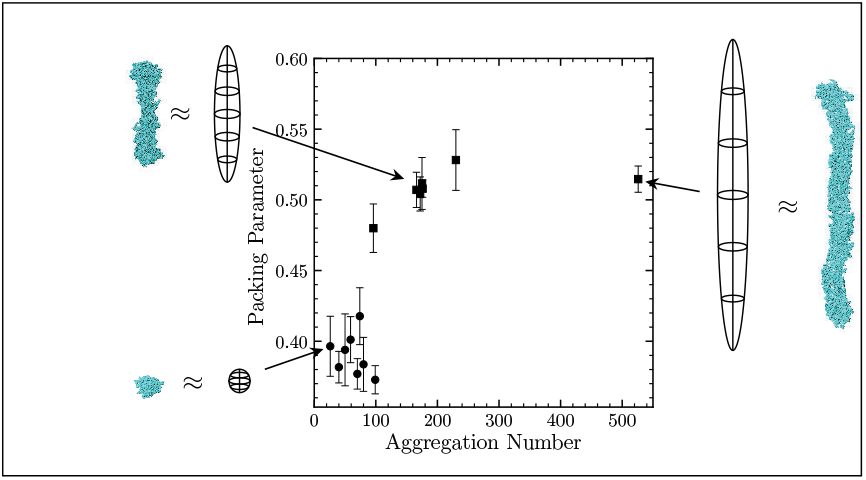

## Introduction

Peptide amphiphiles (PAs) are a class of biomimetic materials traditionally comprised of a hydrophobic moiety connected to a hydrophilic peptide. This modular structure not only allows for the tuning of both the mechanical properties and bioactivity of the resulting self-assembled structures, but it also allows PAs to be useful for a number of potential biomedical applications, including cancer therapy,^1–3^ regenerative medicine,^4,5^ and vaccination.^6–8^ Akin to traditional “surface active agents” (surfactants), such as sodium dodecyl sulfate, the amphipathic nature of PAs facilitates their self-assembly in solution, leading to the formation of peptide amphiphile micelles (PAMs).^9,10^ Like traditional surfactant micelles, PAMs can be generated in a variety of sizes and shapes, depending on their chemical structure^6^ and the conditions of the solution in which they reside (*e*.*g*., total amphiphile concentration, salt concentration,^11^ pH,^4,12^ and temperature).

The size and shape distributions of micelles in solution have a considerable influence on their thermodynamic, transport, and therapeutic properties. Just as the viscosity of an aqueous surfactant solution consisting of cylindrical micelles has been observed to be significantly higher than that of spherical/globular micellar solutions,^13^ the size and shape of micelles used in nanomedicine have been shown to influence several aspects of their bioactivity. For example, size and shape have been shown to influence PAM efficacy in cellular uptake^14–16^ and targeted drug delivery^17,18^ as well as dramatically influencing their capacity to either elicit^6–8^ or suppress^19^ immune responses.

We seek to leverage the significant effort that has been devoted to characterizing the size and shape distributions of traditional surfactant micelles in aqueous solution to develop a model for predicting these properties for peptide amphiphile micelles. There has also been significant effort made over the years to develop mathematical models of micellization. Tanford’s early work^20–22^ on micelle formation put forward an empirical description of the phenomenon wherein the finite size of self-assembled aggregates was the result of a balance between attractive hydrophobic and repulsive electrostatic forces. Israelachvili *et al*.^23^ built upon this framework, incorporating molecular packing within the aggregate core to develop a geometry-based approach to predict the formation of different aggregate shapes. With these changes, the formation of spherical and cylindrical micelles and spherical bilayer vesicles, as well as transitions between these structures, could be explained on the basis of general thermodynamic principles, Tanford’s free energy model, and the geometric constraints imposed by molecular packing considerations. A few years later, Nagarajan and Ruckenstein’s statistical-mechanics–based approach^24,25^ shed light on the physical origin of the attractive and repulsive components of the free energy of micellization and posited steric repulsion between head groups, interfacial tension, and reductions in translational and rotational degrees of freedom as sources of repulsive forces that explain finite nonionic micelle formation.

Analogous to the factoring of the partition function in their earlier model, Nagarajan and Ruckenstein^26^ developed one of the earliest, strictly *a priori* models of aggregation via a molecular thermodynamic (MT) approach. In an MT model, the free energy change associated with the formation of the surfactant aggregate is expressed as the sum of several free energy contributions, all of which can be computed from the chemical structures of the micellar components. MT models of micellization have since become increasingly complex and have permitted the modeling of systems ranging from non-ionic surfactants^27^ to ionic and zwitterionic surfactant systems,^26,28^ binary surfactant mixtures,^28,29^ and even ternary mixtures using regular solution theory.^30^ However, the success of these techniques was largely confined to explaining the aggregation behavior of simple surfactants with small head groups and linear hydrocarbon tails. In light of this observation, some researchers have turned to computer simulation to probe the complex interactions involved in micellization.^31^

In theory, molecular dynamics (MD) simulations based on an atomistic potential energy model have the ability to model arbitrarily complex chemical structures and permit quantitative predictions of micellization behavior. These aspects are especially attractive for the study of PAMs due to the variety and complexity of the interactions they can undergo. In addition to the interactions typically associated with micellization (*e*.*g*., hydrophobic, electrostatic, and interfacial interactions), the peptide headgroup in PAMs can participate in many of the same interactions involved in protein–protein interactions (*e*.*g*., hydrogen bonding, specific inter amino-acid interactions, salt bridge formation, and disulfide bonding).^10^ Unfortunately, these simulations are limited by the computational requirements needed to simulate the sizes and densities of micellar systems,^32^ and currently, all-atom detail can only be resolved at concentrations well above the critical micelle concentration (CMC). To combat the computational requirements necessary to observe micellization in MD simulations, Lee *et al*.^33^ began their simulation of palmitoylated SLSLAAAEIKVAV PAs from a seeded cylindrical configuration and showed stabilization of the cylinder radius within 40 ns. This seeding procedure was then employed in MD simulations applying an external bias to uncover mechanistic insights into the early stages of PA self-assembly^34^ and quantitative estimates for the free energy changes in the assembly process.^35^

While these simulations have uncovered interesting atom-level interactions that lead to the formation of PAMs and even provided quantitative free energy estimates, the relative importance of the various contributions (*e*.*g*., the hydrophobic effect versus tail packing free energy) outlined in the MT modeling approach to micellization are obscured, and it is difficult to extrapolate the results from one amphiphile to another. Stephenson *et al*. published a series of articles^36–38^ outlining a “computer simulation, molecular-thermodynamic” model that addresses this issue in non-PA surfactants. While this framework was originally designed to model amphiphiles whose head and tail groups are ill-defined under the MT convention, we seek to extend this model to PAMs. The first step in this endeavor is to establish a consistent, reproducible means of characterizing the shape of simulated micelles so that shape-dependent free energy contributions such as the tail packing free energy can be appropriately handled. Interested readers are referred to several reviews^13,39,40^ that provide a comprehensive history of modeling the behavior of self-assembling surfactants.

In this work, we develop a quantitative means for shape classification from MD simulations consistent with the original measures developed by Israelachvili.^23^ This shape quantification model is then applied to the estimation of a shape-dependent packing free energy contribution and is show to be consistent with both empirical and theoretical results. The outlined mathematical framework allows the general quantification of the shape of a micelle and the free energy penalty associated with packing the aliphatic core into that morphology for use in the context of a molecular thermodynamic model.

## Theory

### Previous Shape Classification Schemes

The influence PAM shape has on in its capacity to illicit high IgG titers in mice makes predicting PAM morphology important for screening vaccine candidates before investing in experimental research.^6,7,14^ While the atomic resolution of molecular dynamics (MD) simulations is helpful in understanding the relative importance of the various components of PAMs, such simulations are intractable for large collections of molecules. It is thus necessary to devise a method for classifying the morphology of a simulated PAM without atom-scale resolution.

Historically,^23^ morphology classification has been defined on the basis of a dimensionless geometric packing parameter called the “critical packing parameter” or “local packing parameter,” denoted here by *p*. This parameter was originally^23^ defined as the ratio of the volume of the hydrocarbon chain of an amphiphile, *v*, to the product of its optimal surface area, *a*_0_, and a critical chain length 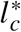, *viz*.,

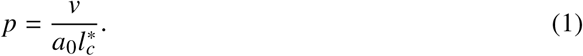

The critical chain length is a semi-empirical parameter that sets a limit on how far chains can extend, though shorter extensions beyond this are allowed. For ideal or “limiting shapes,”^41^ the critical chain length corresponds to a maximum for a dimension of the shape from which the total volume and surface area of the micelle core can be calculated. For example, in spheres 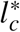 represents an upper bound for the radius of the core of a spherical micelle, 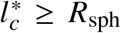, and the total volume and surface area of the core follow from geometry as 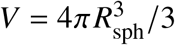 and 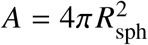, respectively.

Then, the overall packing parameter or simply, the packing parameter, *P*, for this limiting shape can be estimated via,

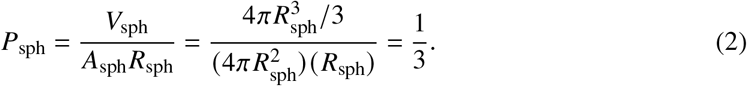

If each of the composite quantities in Equation (2) is narrowly distributed about its mean value, *p* in a micelle comprised of *g*_*k*_ amphiphiles for an individual amphiphile can be recovered,

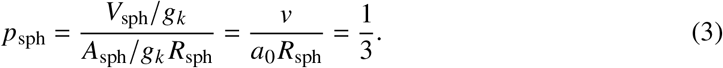

This approximation is certainly reasonable in limiting morphologies where the curvature of the shape is uniform throughout. Using a similar approach, *p* can be used to define critical conditions for infinite cylinders and planar bilayers at values of 1/2 and 1, respectively. These limiting shapes delineate regions of *p* (and *P*) that can be used to classify the shape of the core of the micelle in the vicinity of an amphiphile for which *p* is estimated. For general amphiphiles, *v* is taken to be the volume of the hydrophilic portion, *a*_0_ becomes the surface area of the amphiphile’s equilibrium hydrophilic interface,^42^ *a*, and 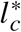, the critical extension length, becomes the extension of the hydrophobic chain, *l*_*c*_. Upon estimation of these quantities, the local packing parameter can be calculated, *p* = *v*/(*al*_*c*_), where the shape of the core can be classified as follows: *p* ≤ 1/3 for spherical, 1/3 ≤ *p* ≤ 1/2 for ellipsoidal, *p* ≈ 1/2 for rodlike, 1/2 ≤ *p* ≤ 1 for interconnected structures, and *p* ≈ 1 for vesicles and extended bilayers.

While generally helpful, nanoparticles and micelles are often observed to be in morphologies outside of these ideal shapes. Unfortunately, in these conformations, the local packing parameter of an individual amphiphile is not always representative of the geometry of the entire core. On the other hand, the total packing parameter depends on geometric properties of the entire aggregate core. However, without assuming that the shape of the particle is one of these ideal morphologies, the interpretation of *l*_*c*_ in the context of the area and volume of the micelle core is ambiguous, causing there to no longer be a succinct form that can be used to calculate *P*. Furthermore, for a collection of point masses as obtained from an MD simulation, the concepts of volume and equilibrium surface area are not well-defined, and quantifying them consistently is not straightforward.

Since the original classification system was defined, several new measures for quantifying the shape of polymeric polypropylene chains based on the principal axes of gyration have been developed^43,44^ and applied in their native form to characterize the shape of simulated nonionic surfactant micelles.^45^ In these studies, the asphericity, acylindricity, and relative shape anisotropy indices were related to expected symmetry relations of the gyration tensor to define measures that obey specific limiting behaviors. For example, the relative shape anisotropy can assume values between 0 and 1, corresponding to shapes with at least tetrahedral symmetry and a “linear array of skeletal atoms,” respectively. In this work, these two shape assignment schemes are bridged: the relatively ambiguous, macroscopic shape assignment scheme associated with the packing parameter is quantified via the assumption of an underlying shape model whose principal moments of inertia can be determined by the masses and coordinates of the particles comprising the micelle. First, an effective ellipsoid is described by the lengths of three axes (*a, b, c*) defined by the eigenvalues of the moment of inertia tensor obtained from a collection of point masses. Next, a dimensionless index defined by the eigenvalues of the moment of inertia tensor is defined to provide a quantitative metric for the shape. Then, to ensure that morphologies classified by this index and the packing parameter are consistent, the definition of the packing parameter (which was previously restricted to idealized geometries) was expanded via the introduction of a “characteristic length.” Next, the ellipsoidal shape model was employed to generalize a packing free energy model as a smooth function of the shape and size of the micelle core. Finally, these measures were tested for how well they agree with the literature and obey expected scaling relationships or phenomenological models.

### Extension to Intermediate Morphologies

The major, intermediate, and minor axes of a uniform ellipsoid (*a, b*, and *c*, respectively) can be uniquely defined by the diagonal (body-frame) moment of inertia (MOI) tensor obtained from the simulation,

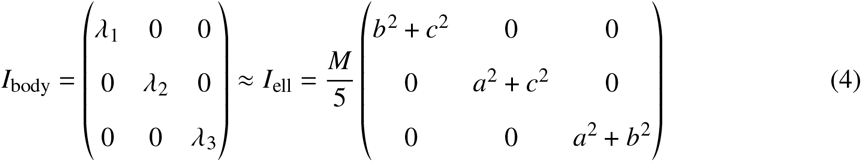

where *λ*_*i*_ is the *i*^th^ eigenvalue of the MOI tensor, which is given in terms of the radius of gyration about the *i*^th^ axis, *R*_g,*i*_, and total mass, *M*, as 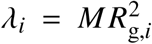. We have adopted the convention *λ*_1_ ≤ *λ*_2_ ≤ *λ*_3_, which is equivalent to *a* ≥ *b* ≥ *c* in terms of the dimensions of the best-fit ellipsoid. Another popular definition from the polymer community, referred to here as the “IUPAP definition,” uses a single radius of gyration, *R*_g_ which is related to the definition of the radii of gyration about the principal axes via 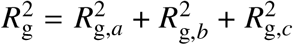. For a collection of point masses, this tensor (in the lab frame) is given by,

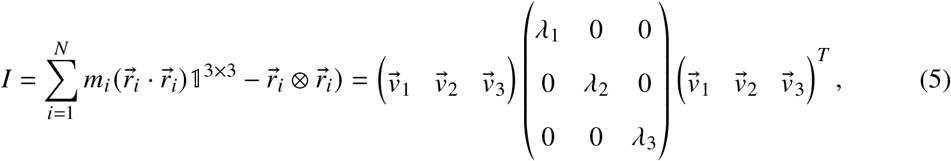

where *m*_*i*_ is the mass of the *i*^th^ particle, 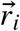 is the vector pointing from the center-of-mass of the collection of point masses to the position of the *i*^th^ particle, ⊗ indicates the outer product, 𝟙^3×3^ is the 3 × 3 identity matrix, *N* is the total number of point masses comprising the micelle, and 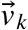 is the *k*^th^ principal axis vector.

Following some algebraic manipulation, the axes of the model ellipsoid can be obtained in terms of the radii of gyration of the collection of point masses in the simulation about their principal axes, as follows:

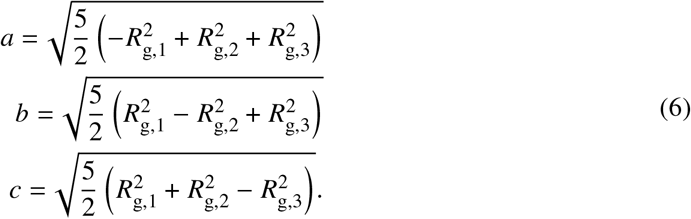

In this fashion, we have assumed the point mass distribution can reasonably be captured by an ellipsoidal shape model and uniquely defined by its parameters. It should be noted however, that while the ellipsoid shape model permits a smooth and unique quantification of the shape of a collection of point particles, this approach inherently produces a shape that exhibits central symmetry, irrespective of whether that assumption is consistent with the provided mass distribution.

In addition to defining a model ellipsoid uniquely, the MD-derived moment of inertia tensor was leveraged to define an index that captures the shape. This index, termed the “normalized relative shape anisotropy,” 𝒜,

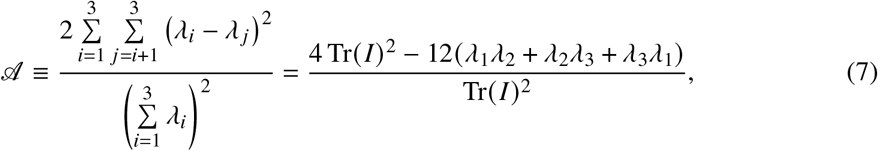

whose right-hand side can be defined from two of the three invariants of *I*, has a value of 0 for a body with at least tetrahedral symmetry (*e*.*g*., a perfect sphere), and approaches 1 for a cylinder of infinite length. In accordance with its name, the index is simply four times the relative shape anisotropy,^43,44^ allowing the planar symmetric infinite cylinder to have 𝒜 = 1 rather than a configuration wherein all points are co-linear. The advantages of such an index are that it not only provides a quantitative means for characterizing morphologies between a perfect sphere or infinite cylinder, but that it can also be estimated without an *a priori* assumption of the morphology’s symmetry nor prescription of a shape model.

Once a shape model has been deemed appropriate, however, 𝒜 can be expressed on a dimensionless basis in terms of scale-invariant dimensions, or aspect ratios, of the assumed shape. For example, in an ellipsoid, these aspect ratios are taken to be *α*_1_ = *b*/*a* and *α*_2_ = *c*/*a* so that they are confined to the intervals (0, 1) and (0, *α*_1_), respectively. In terms of these reduced dimensions (*i*.*e*., the aspect ratios of an ellipsoid), 𝒜_ell_ is

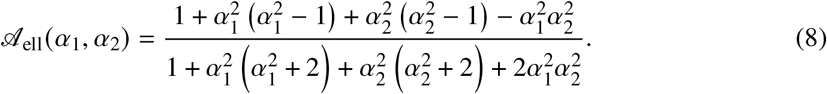

The power of this scale-invariant definition will become evident for distributions that exhibit high symmetry. Figure 1 provides a frame of reference for the aspect ratios defined for a general ellipsoid. Contours are provided to help visualize the scaled relative shape anisotropy as well as an asymmetry threshold, 𝒯, between the two largest principal moments of inertia (*i*.*e*., the eigenvalues of the inertia tensor) from which the ellipsoids are fit,

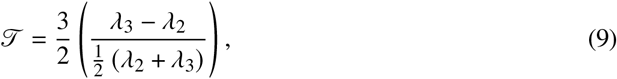

corresponding to a ratio between the difference in the moments of inertia about the two minor axes and their mean values. This index is then scaled by a factor of 3/2 so that the range of the asymmetry threshold spans the interval [0, 1], corresponding to *α*_1_ = *α*_2_(for which : 𝒯 = 0) and (*α*_1_, *α*_2_) = (1, 0) (for which : 𝒯 = 1), respectively. Previously,^46^ a direct ratio of these principal moments of inertia was used as means to assess asymmetry,

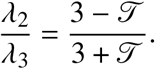

While both measures 𝒯 and *λ*_2_/*λ*_3_ are monotonic with the increase in symmetry, the convenient interval associated with 𝒯 and the highly symmetric bodies observed in our simulations motivate its use in this work.

**Figure 1:**
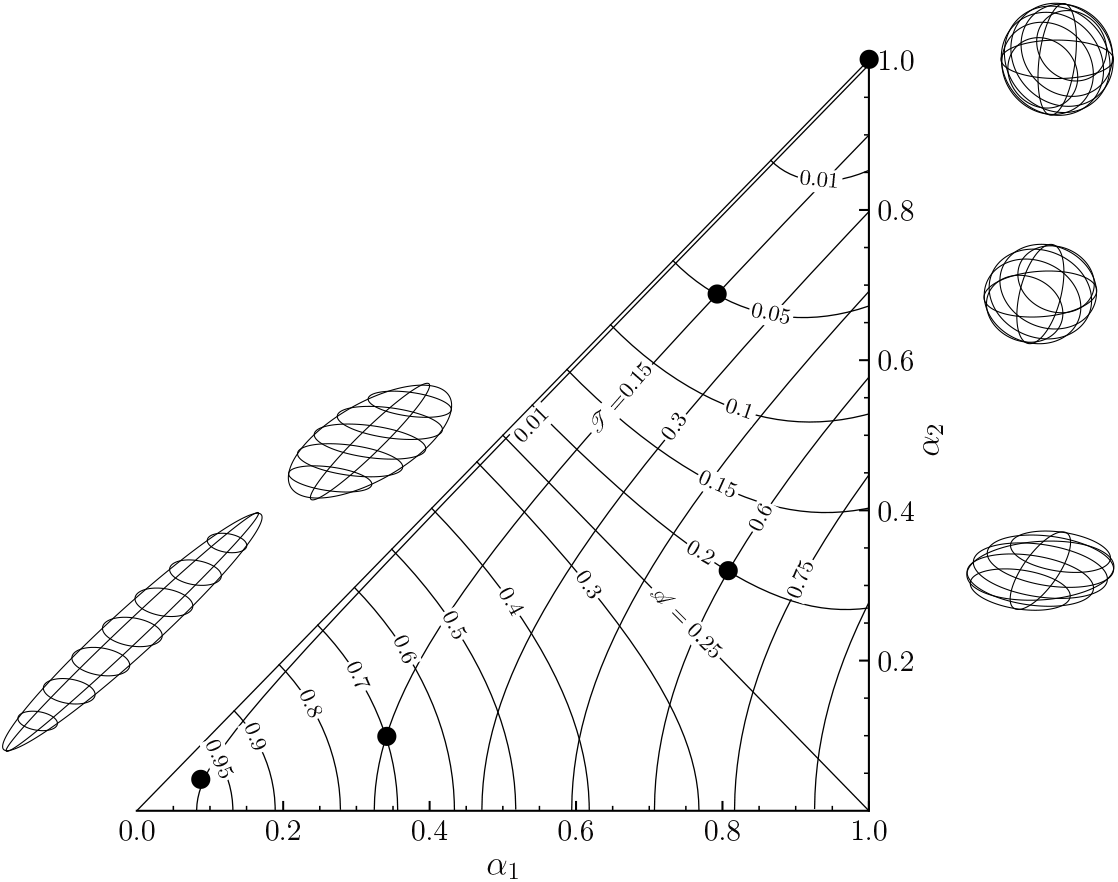
Morphology diagram for general ellipsoids with major aspect ratio *a*_1_ and minor aspect ratio *a*_2_ where the depicted ellipsoids share the same volume. Contours indicate constant values of the normalized relative shape anisotropy index 𝒜 (Equation 8) or constant symmetry threshold : 𝒯 (Equation 9). Indicated points correspond to the aspect ratios of the depicted ellipsoids.

In Figure 1, as both *α*_1_ and *α*_2_ approach 1, the ellipsoid becomes increasingly spherical, and the value of the normalized relative shape anisotropy approaches zero. When *α*_1_ decreases at constant *α*_2_, the ellipsoid becomes increasingly prolate or “stretched out” along the main axis whereas, a “flattening” or oblate behavior better describes the decrease in *α*_2_ at constant *α*_1_. Along constant 𝒯 contours, ellipsoids change from a morphology resembling a baguette at low (*α*_1_, *α*_2_) to one hardly distinguishable from a sphere. Interestingly, 𝒜 contours make a transition from concave up to concave down as this index increases through 𝒜 = 0.25—the value approached by an infinite lamella (*α*_1_ → 1^−^, *α*_2_ → 0^+^).

While a best-fit ellipsoid is a natural choice for morphologies with three unique radii of gyration, the choice of the shape model fit to the calculated MOI tensor becomes more ambiguous as the symmetry of the body increases. For example, for a system with *α*_1_ < *α*_2_ = *α*_3_, unique dimensions for a capsule (*i*.*e*., a cylinder with two hemispherical end-caps of the same radius), prolate spheroid (*i*.*e*., an ellipsoid with *b* = *c*), or cylinder can all be defined. Since each of these shape models is derived from the same MOI tensor, they share the same 𝒜 value but not necessarily the same aspect ratio. Figure 2 illustrates the dependence of 𝒜 on the aspect ratios of several shape models with expressions summarized in Table 1. By design, the normalized relative shape anisotropy of each of the shape models that exhibits radial symmetry about its primary axis (*i*.*e*., finite cylinders, prolate ellipsoids, and capsules) approaches that of an infinite cylinder as the aspect ratio goes to 0, *viz*.,

**Table 1:**
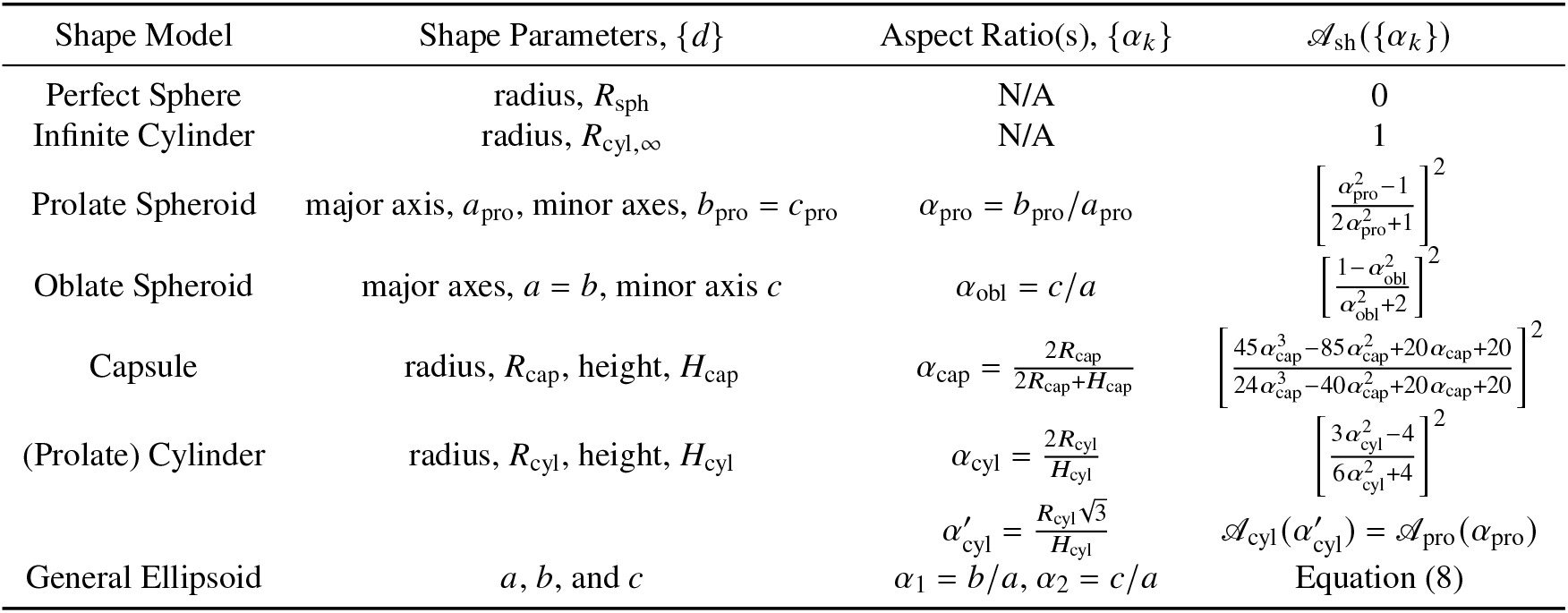
Expressions for the parameters, aspect ratios, and normalized relative shape anisotropy index, 𝒜_sh_, for different shape models.

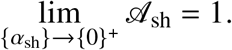

However,

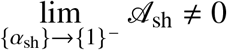

for each shape model, sh, in this work. Specifically, 𝒜_cyl_, the normalized relative shape anisotropy of a prolate cylinder, does not equal zero as its aspect ratio goes to 1 and instead, it achieves a value equal to that of a sphere when 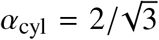. Interestingly, if this factor is used to redefine the aspect ratio of a prolate cylinder,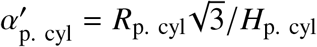, the dependence of the normalized relative shape anisotropy index on *α*′_p. cyl_ is identical to that of a prolate spheroid. The limiting behavior of an oblate spheroid is also interesting, as the constraints from the definitions of the axes (*i*.*e*., *a* ≥ *b* ≥ *c*) and the requirement that *α*_1_ = 1 limit 𝒜 to a maximum value of 1/4. This limit, taken to be representative of an infinite plane, is attained as *a* = *b* ≫ *c*, and the approach can be visualized by the compression of a sphere between two parallel planes as *α*_1_ goes from 1 to 0.

**Figure 2:**
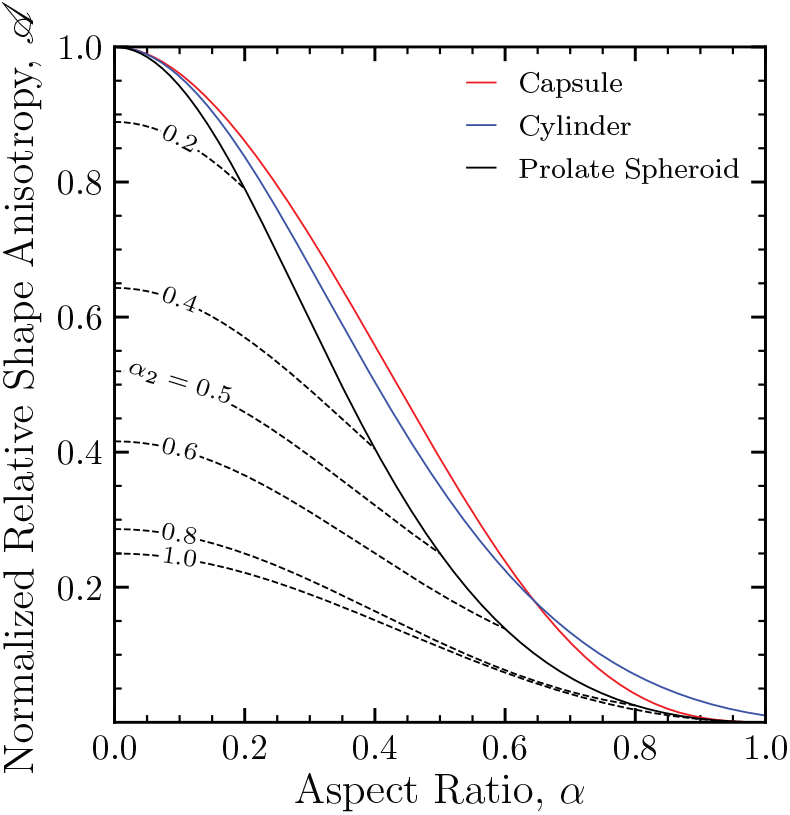
Normalized relative shape anisotropy, 𝒜, as a function of aspect ratio, *α*, for several shape models. Solid lines indicate shape models whose primary axis exhibits rotational symmetry (black, prolate spheroid; red, capsule; blue, prolate cylinder), whereas dashed lines are used for general ellipsoids with *α*_1_ = 1 but varying *α*_2_ values corresponding to an oblate spheroid. If the aspect ratio of the prolate cylinder is redefined from the ratio of the largest width to the largest height so that 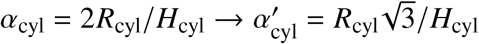, the corresponding curve coincides with the prolate spheroid.

For morphologies that are symmetric about the primary axis, the specification of the scaled relative shape anisotropy index is sufficient to fix the relative proportion of geometric properties (*i*.*e*., *V*, *A, l*_*c*_, or *P*) of the two shape models. For example, the ratio of the surface area of a capsule to that of a prolate spheroid is uniquely determined by *α*_1_ and *α*_2_. Equating the smallest eigenvalues of the prolate spheroid and capsule results in an expression for *R*_cap_ in terms of *b*_pro_,

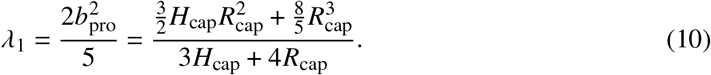

Then, rearranging the expression for the aspect ratio of a capsule to get an expression for *H*_cap_ and requiring *α*_cap_ ≠ 0,

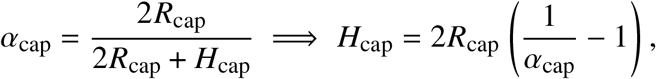

which results in

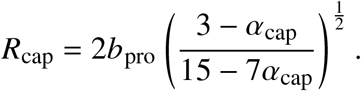

Recall that *α*_cap_ lies in the interval (0, 1) and *R*_cap_ > 0. The ratio of the area of the capsule to that of prolate spheroid with the same eigenvalues of the moment of inertia tensor can be expressed in terms of their aspect ratios,

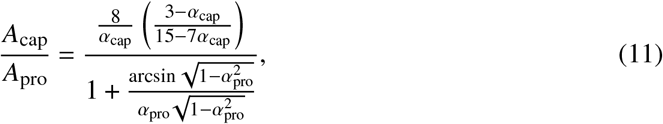

which are uniquely determined by the value of 𝒜 according to the expressions in Table 1. These facets provide an opportunity to visualize this relatively abstract index (Figure 3). For shapes fit to identical (symmetric) moment of inertia tensors, reasonable agreement (ratios near 1) in the volume and area ratios between the spheroid and both the capsule and the cylinder is observed. This suggests that if a body described by a moment of inertia tensor was modeled as a prolate spheroid, when a cylinder or capsule would likely better capture the morphology of the body, only a modest error would be incurred. When modeling the body as a prolate spheroid, a systematically larger estimation of both the surface area and the volume relative to the capsule shape model is observed. Alternatively, the best-fit prolate spheroid has regions of both larger and smaller surface areas than the corresponding best-fit prolate cylinder. The ratio of the volumes of these two shape models is constant with 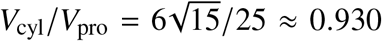. Indeed, the most egregious difference in the surface area estimate between the prolate spheroid and capsule shape models occurs in bodies with 𝒜 ≈ 0.764, for which *A*_cap_/*A*_pro_ ≈ 0.821, fand the largest discrepancy in the volume estimated by these two models occurs in bodies with 𝒜 ≈ 0.629 wherein *V*_cap_/*V*_pro_ ≈ 0.814. These scenarios amount to 21.8% and 23.0% larger estimates of the surface areas and volumes, respectively, when comparing the prolate spheroid to the capsule shape model.

**Figure 3:**
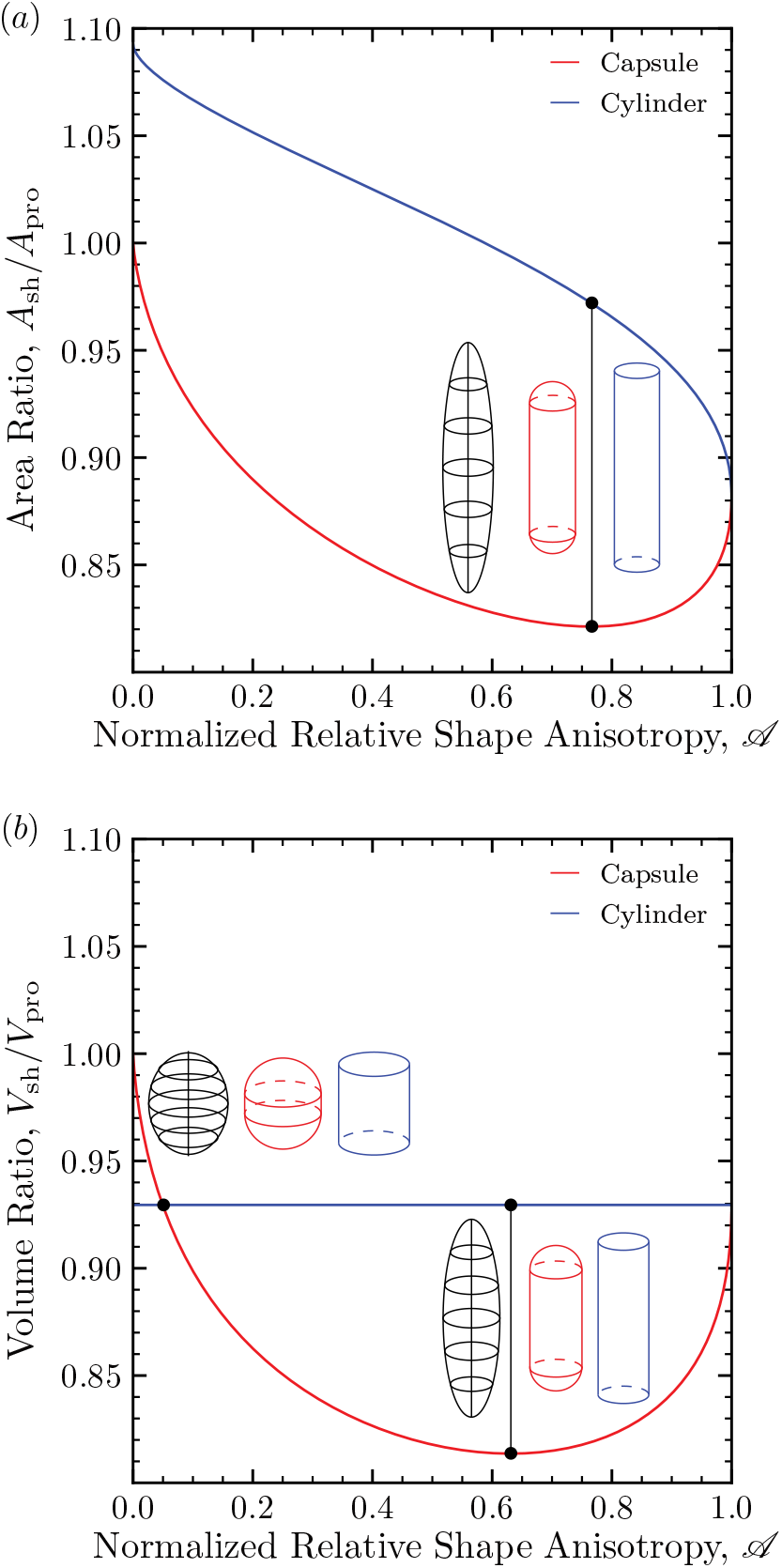
Parameter analysis of a radially symmetric moment of inertia tensor for cylindrical and capsular shape models. The (a) area and (b) volume are normalized to the corresponding value obtained for the prolate ellipsoid. Representative renderings of the specified shape models and the correspond prolate ellipsoids sharing the same eigenvalues of the MOI tensor are shown for the indicated points.

To ensure morphologies classified by either the normalized relative shape anisotropy index or the overall packing parameter are consistent, the definition of *P* was expanded to capture “intermediate morphologies.” As outlined earlier, the packing parameter is well defined for shapes for which *l*_*c*_ is well defined and becomes ambiguous for “non-ideal”—or “intermediate”—shapes. In their seminal work introducing the critical packing parameter, Israelachvili *et al*.^23^ introduced two methods to extend the applicability of *p* outside of these ideal geometries. The first of these provides thresholds for *p* based on the maximum degree to which the hydrophobic tail can be extended as outlined by the aforementioned classification regions (*e*.*g*., *p* ≤ 1/3 for spherical micelles). This categorical approach has been widely adopted and recently, assuming *p* ≈ *P*, Iakimov *et al*.^47^ successfully applied these classification regions to identify the model for the conformation of a poly(tyrosine) tail that best agreed with the experimental whole aggregate morphologies of poly(L-tyrosine)– PEG block copolymers. The second extension to *p* has gained far less traction.^23^ This approach introduces an approximate expression for *p* based on three geometric parameters: the length of the hydrocarbon region of the amphiphile and two radii of curvature. While this approach permits the estimation of *p* for a wide class of morphologies, it remains rather abstract, as the estimation of the radii of curvature remain challenging.

Here, we generalize the definition of *P* by introducing a procedure for defining the characteristic length for an assumed shape model. That is, the overall packing parameter is estimated after assuming a shape model via known expressions for the volume and area of the chosen shape, as well as an aspect ratio (α_sh_) and scale-dependent characteristic length (*l*_c,sh_), *viz*.,

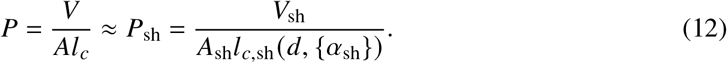

In this fashion, the *P*_sh_ as described here no longer provides thresholds for idealized shapes and instead can be smoothly estimated for real shapes after the assignment of a shape model.

Expressing 𝒜 in terms of aspect ratios and introducing an aspect-ratio–dependent characteristic length provides an avenue for ensuring the two classification schemes are consistent with one another for idealized shapes. For simplicity and consistency, the characteristic length for a given shape model was taken to be the lowest-degree polynomial in which agreement could be observed: values of the aspect ratios that would assign a given shape as a sphere, infinite cylinder, or plane via 𝒜_sh_ are the same as the aspect ratios that would classify the morphology as a sphere, infinite cylinder, or plane via *P*_sh_. Explicitly, *l*_*c*_ is defined to satisfy the following,

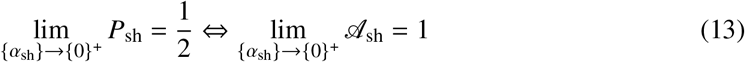

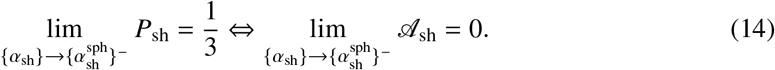

where 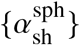 indicates the set of α_sh_ values for which 𝒜_sh_ of that shape model has a value of 0. For example, for a capsule, 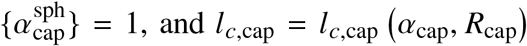 is found as follows. First, the packing parameter of a capsule, *P*_cap_, is expressed as a function of the aspect ratio, α_cap_, and temporarily, the parameter, *R*_cap_,

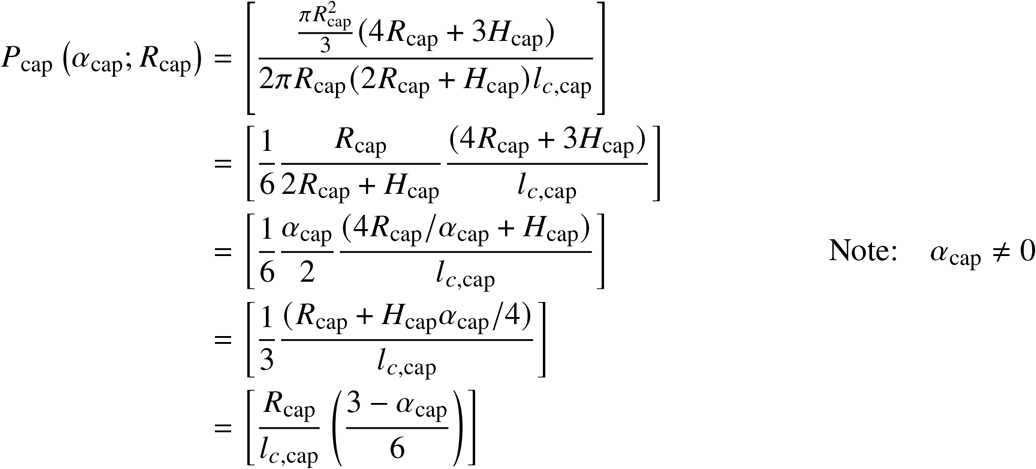

In the infinite cylinder limit,

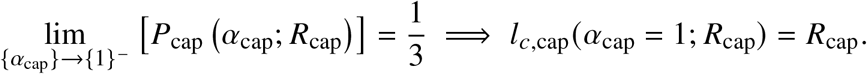

Likewise, for a perfect sphere,

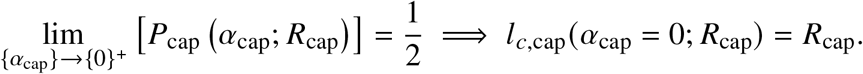

Finally, the lowest-degree polynomial that satisfies both of these conditions is clearly

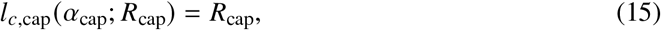

as no *α*_cap_ dependence is necessary to achieve the desired limiting behavior. Expressions for the characteristic lengths of other shape models are found similarly and are summarized in Table 2. Furthermore, as expected for a dimensionless index, the packing parameter can be shown to be independent of the absolute scale of the capsule, as its dependence on the dimension *R*_cap_ vanishes upon substitution of the expression for the characteristic length, *viz*.,

**Table 2:**
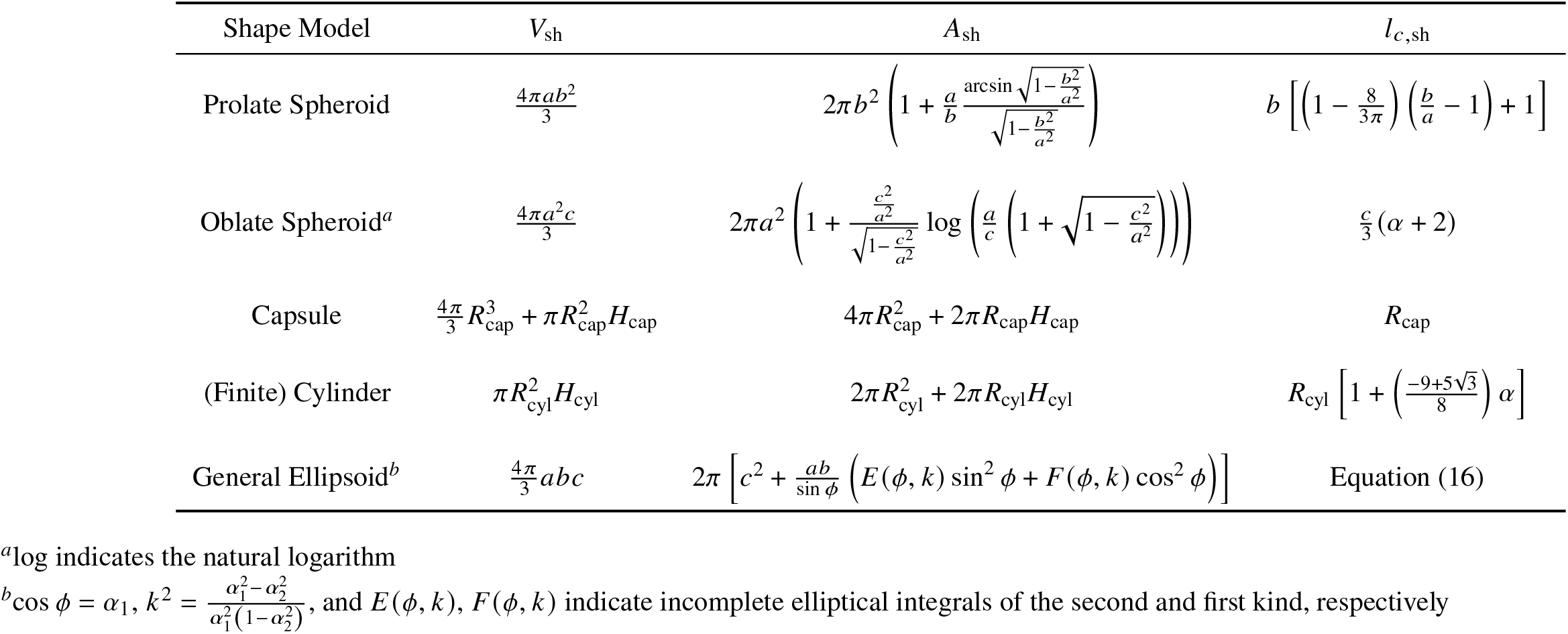
Expressions for the volume, surface area, and characteristic lengths of select shape models.

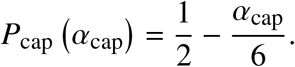

While the aforementioned conditions are sufficient for radially symmetric objects, the characteristic length of a general ellipsoid is also required to be equivalent to that of a prolate spheroid or that of an oblate spheroid in the appropriate symmetry limits. The lowest degree polynomial consistent with these expressions is found to be a quadric surface whose coefficients are obtained via the Moore–Penrose inverse^48^ of the system of equations implied by the functional form,

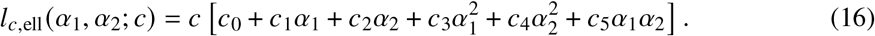

The coefficients in Equation 16 are 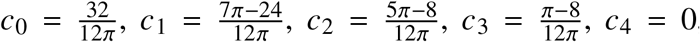, and 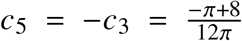. The hyperbolic paraboloid de(scribed by su) ch a surface is rotated about the *l*_*c*_-axis by π/8 and has its origin 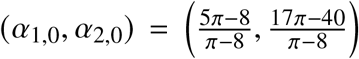 outside the region of interest (0 ≤ *α*_1_ ≤ 1) ∪ (0 ≤ *α*_2_ ≤ *α*_1_).

Similar to the assessment of the surface area and the volume of the best-fit ellipsoid, these generalizations of the characteristic length and packing parameter are compared for prolate ellipsoids, capsules, and cylinders in Figure 4. The characteristic length of the capsule shape model is systematically larger than that of the prolate ellipsoid with the same moments of inertia, with the largest discrepancy achieved in the limit 𝒜 → 1, resulting in a 5.1% smaller estimate of the characteristic length by the prolate spheroid compared to either the capsule or cylinder models. Interestingly, the prolate spheroid makes a transition from under-to overestimation of the characteristic length of a finite cylinder, and the largest difference is seen in the cylindrical limit. The normalized relative shape anisotropy at which *l*_*c*,cyl_ = *l*_*c*,pro_ is 𝒜 ≈ 0.726, and representative renderings of the shapes are indicated. These renderings highlight the notion that the characteristic length is not simply the largest radial dimension: despite equivalent characteristic lengths, the minor axis of the spheroid and the radius of the cylinder are different.

**Figure 4:**
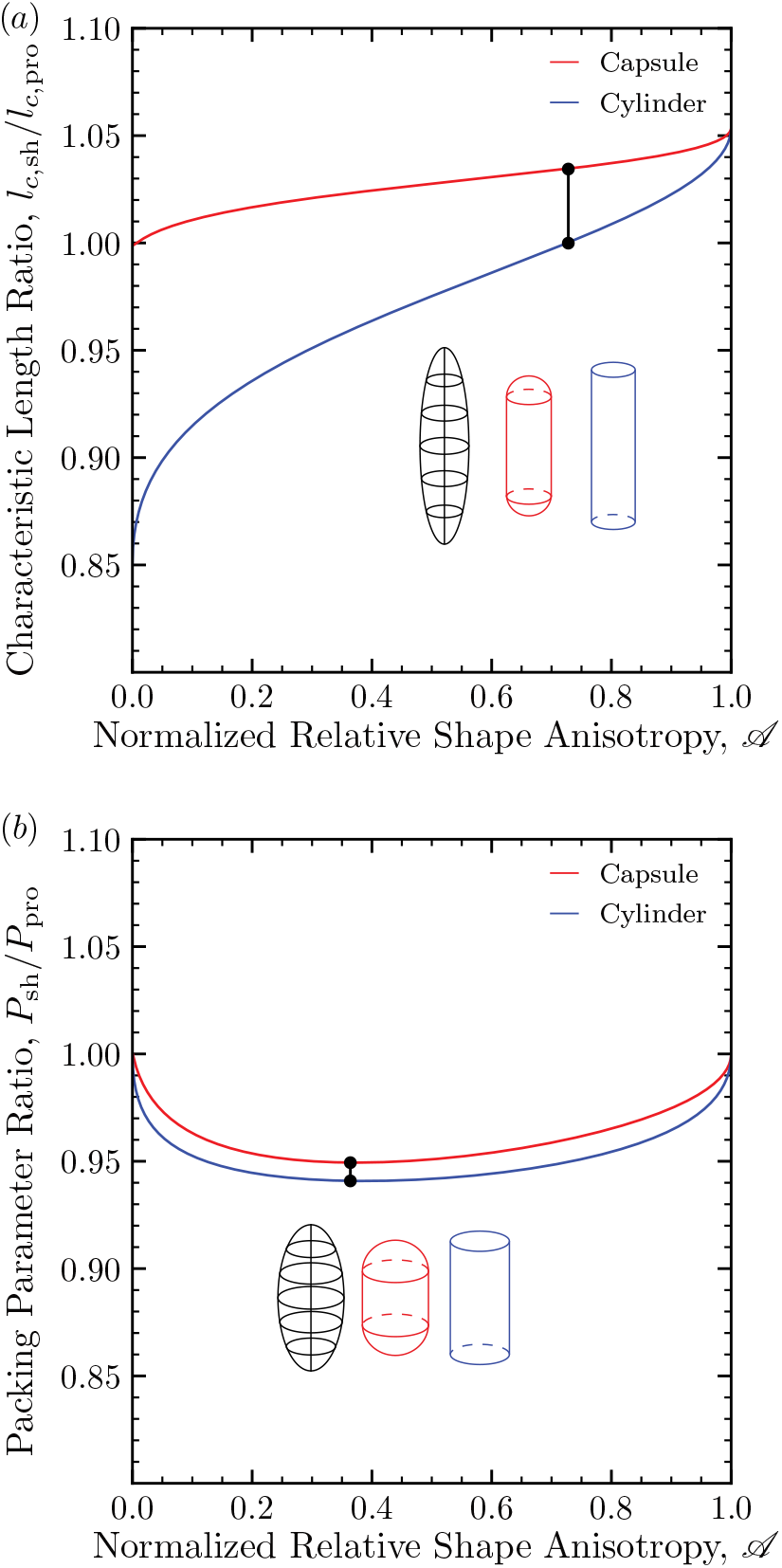
Assessment of the polynomial *l*_*c*_ model and packing parameter model for prolate cylinder and capsular shape models relative to the corresponding value of the prolate spheroid. Represen-tative renderings of the specified shape models and the corres(pond) prolate(ellip)soids sharing the same eigenvalues are shown for (a) *l*_*c*,cyl_ = *l*_*c*,pro_ and (b) min 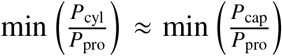. By design, the packing parameter for each shape model approaches the same value for perfect spheres (𝒜 = 0) and infinite cylinders (𝒜 = 1).

The *P* for an ellipsoid is systematically larger than that for both the capsule and the finite cylinder. By design, equality is achieved for perfect spheres and in the infinite cylindrical limit. The largest percentage difference in the packing parameter of the prolate spheroid relative to the capsule and cylinder shape models occurs for bodies with 𝒜 ≈ 0.362 at 5.37% and 6.29% larger, respectively.

### Application to Tail Packing Free Energy

The “tail packing free energy” (TPFE) was first introduced in molecular thermodynamic models of micellization as an empirical correction to explain the discrepancies in approximating the hydrophobic core as a bulk hydrocarbon phase when modeling the free energy change incurred when the hydrophobic tails of the amphiphiles move from an aqueous environment to the hydrophobic core.^21^ Models for the TPFE of micellization approximate the energetic penalty associated with the geometric packing constraints that arise inside the core of the aggregate that is otherwise absent in a bulk hydrophobic phase. This contribution to the standard free energy change of micellization, 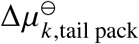, has since been shown^49–52^ to be dependent on both the size and shape of the hydrophobic core. Of these analyses, the analytical, shape-dependent expressions of Semenov^53^ applied to micelles by Nagarajan *et al*.,^26^ referred to henceforth as the Semenov–Nagarajan model, provide an opportunity to leverage the ellipsoidal shape model to predict the packing free energy for the cores of micelles of intermediate shapes.

Recently, Danov *et al*.^39,54^ have also extended the Semenov–Nagarajan model. In their approach, called the “Danov Model” (DM), a unified expression for the per-amphiphile packing free energy is derived by the introduction of a unified, local-packing-parameter–dependent form of the packing constraints of the Semenov–Nagarajan model. In this fashion, the DM permits the estimation of the packing free energy per amphiphile for an amphiphile in a region (such as the spherical endcaps of a spherocylinder) of local packing parameter, *p*,^39^

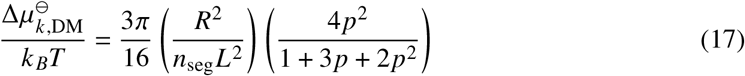

where *R* is well-defined for spheres and infinite cylinders. Unfortunately, surfaces of constant segment density within the core of ellipsoidal micelles, such as those presented in this work, cannot readily be put in the form of the DM boundary conditions. The model presented here shares the same limiting behavior as the DM model with the added benefit of extensibility to additional shape models.

Following the derivation of the Semenov–Nagarajan model, we estimate the total packing free energy of the hydrophobic core for three limiting cases. For a perfectly spherical core, 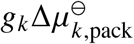 is modeled^26^ by

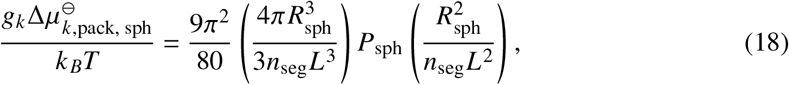

where *P*_sph_ is the packing parameter for a sphere, *R*_sph_ is the radius of the sphere, and *n*_seg_ and *L* are parameters describing the number of segments of the surfactant chain and the size of the lattice, respectively. These were taken to be *n*_seg_ = (*n*_*c*_ + 1)/3.6 and *L* = 0.46 nm, inherited from the Nagarajan’s application^26^ of the Semenov model.^53^ Similarly, the packing free energy of the hydrophobic core per unit length for an infinite cylinder and per unit area for an infinite lamella are given by

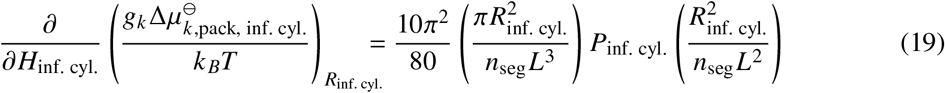

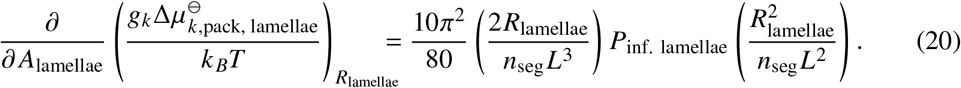

These boundary conditions suggest a model of the form

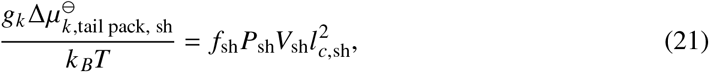

where *f*_sh_ is a shape-model-dependent function introduced to satisfy the boundary conditions. For example, for an ellipsoid, *f*_ell_ = *f*_ell_(*a*_1_, *a*_2_). Since these conditions depend on the shape model assumed in their derivation, we attempt to account for this dependence by assuming that the change in the free energy change of tail packing with respect to the volume is approximately the same for the ellipsoidal model posed here and the shape model used to arrive at the preceding expressions. In the infinite cylinder limit,

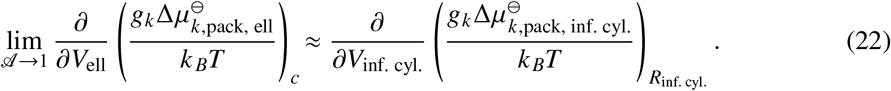

We then apply the chain rule to relate the partial derivative with respect to volume to that with respect to the height of the infinite cylinder,

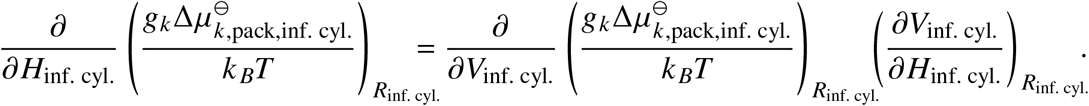

Finally, the limiting case of the infinite cylinder requires the general ellipsoid model to satisfy,

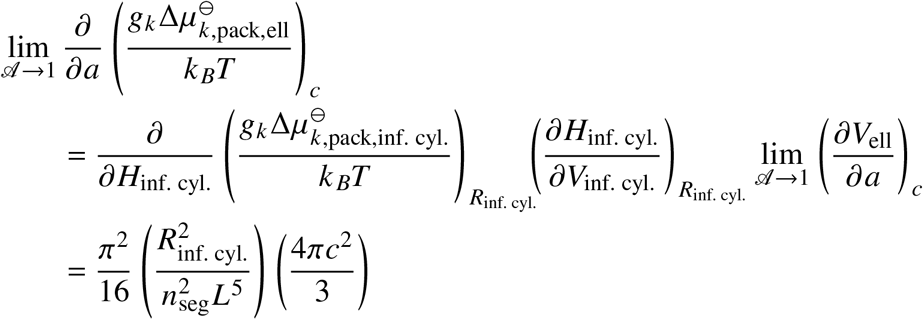

Then, *R*_inf. cyl._ is found in terms of *c* via

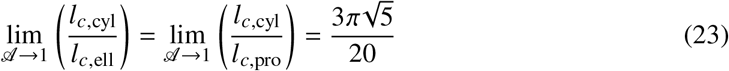

where the ratio *l*_*c*,cyl_/*l*_*c*,pro_ is evaluated by expressing each characteristic length in terms of the two unique eigenvalues of the moment of inertia tensor,

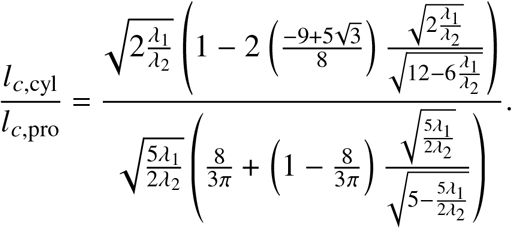

Taking the limit as 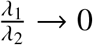 (for which 𝒜 → 1),

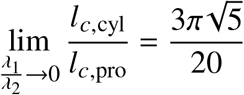

and finally,

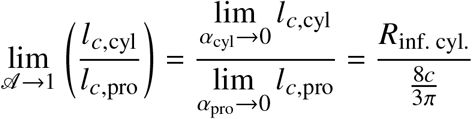

Yielding

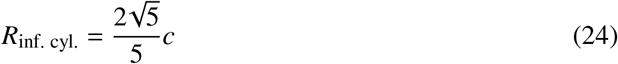

and

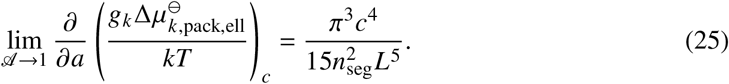

The boundary conditions for the ellipsoid packing free energy model provided by the sphere and infinite lamella cases follow from similar analyses, with results summarized in Table 3.

**Table 3:** Summary of boundary conditions for estimating the contribution of tail packing to the free energy of micellization.

| Shape | Length Relation | Boundary Condition |
| --- | --- | --- |
| Sphere | $R_{\text{sph}} = c$ | $\lim_{\mathcal{A} \rightarrow 0} \frac{g_k \Delta \mu_{k,\text{pack,ell}}^{\ominus}}{k_B T} = \frac{4\pi^3}{80} \frac{c^5}{n_{\text{seg}}^2 L^5}$ |
| Infinite Cylinder | Equation (24) | Equation (25) |
| Infinite Lamella | $R_{\text{lamella}} = \frac{\sqrt{15}}{5} c$ | $\lim_{(\alpha_1, \alpha_2) \rightarrow (1,0)} \frac{\partial}{\partial V} \left( \frac{g_k \Delta \mu_{k,\text{pack,ell}}^{\ominus}}{k_B T} \right)_c = \frac{3\pi^2}{20} \frac{c^2}{n_{\text{seg}}^2 L^5}$ |

From the assumed functional form,

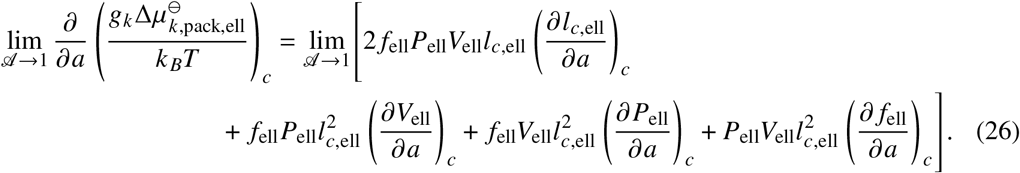

Upon taking the limit, the only surviving term is

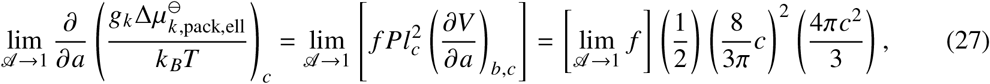

resulting in

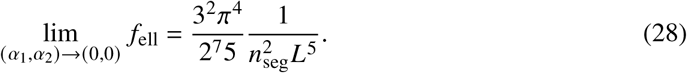

In a similar fashion,

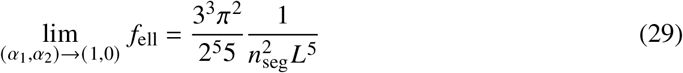

And

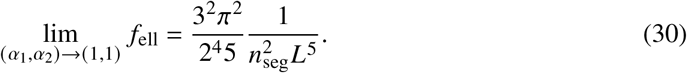

Then, assuming a planar form for *f*_ell_,

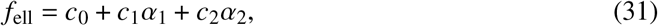

with *c*_0_, *c*_1_, and *c*_2_ are given by the system of linear equations defined by Equations (28) to (31). Finally, assuming an ellipsoidal shape model, the change in free energy associated with packing the hydrophobic tails into the micelle core can be obtained as a smooth function of the shape and size of the core,

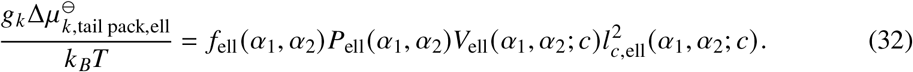

## Methods

### Molecular Dynamics Simulations

Molecular dynamics (MD) simulations were performed using GROMACS^55^ with PAMs seeded in either a spherical or spherocylindrical “capsule-like” geometry with varying aggregation numbers. All solutes (namely peptide amphiphiles and ions) were modeled using bonded and non-bonded interaction parameters from the “CHARMM27” force field^56^ and, accordingly, water was modeled explicitly using the modified TIP3P force field.^56^ Parameters for the pseudo-peptide bond formed upon palmitoylation of lysine were introduced into the force field (see Supporting Information) so that the chemical environment near this synthetic bond mimics the peptide bonds for which the parameters were originally optimized.

The geometry of a di-palmitoylated PA, (palmitoyl)_2_KILRTQSECKEKEKEKE-CONH_2_— Palm_2_K-NA-(KE)_4_ was obtained by connecting a palmitic acid moiety to both of the nitrogens present in a lysine to form pseudo-peptide bonds at both locations. When initially connected, the configurations were those of an extended aliphatic and extended peptide group so as to avoid overlap and artificial entanglement in the combined configuration. This highly contrived structure was then subject to a brief, *in vacuo* minimization to allow the extended conformation to relax. PAM configurations were then seeded according to the procedure described in the next paragraph: Stage 1, simulation set-up; Stage 2, minimization; Stage 3, equilibration; and Stage 4, production run.

First, the configuration of a monomer was oriented such that the “primary vector” (the vector connecting the terminal tail carbon of the aliphatic chain and the alpha carbon of the C-terminal amino acid) was colinear with the *x*-axis. The coordinates of the unimer themselves were slightly offset from the origin (in the +*x* direction) so that subsequent replication did not produce overlapping core atoms.

Second, for micelles initialized as spheres, *g*_*k*_ points (one for each PA) equivalently spaced on the surface of a sphere were generated according to a published method.^57^ The vectors pointing from the origin to each of these points are denoted 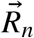. The primary vector of the PA unimer can be oriented to be parallel to 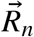 by rotation about the axis perpendicular to both the primary vector and 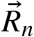 by the angle between the two vectors; thus, a rotation matrix^58^ (for each PA unimer) is applied to each atom in the oriented configuration to generate a “seeded” structure for the spherical PAMs. As a first-order approximation, the Henderson–Hasselbalch^59,60^ equation was then used to assign a protonation state to each amino acid residue in the system as though they were free species in a pH 7.4 solution. That is, no effort was made in the initial geometry to account for the change in the likelihood of a particular protonation state of a side group to account for the local chemical environment (*e*.*g*., another nearby charged side group).

Upon “seeding” the initial micelle structure, the system was solvated via the `gmx solvate` command, which places water molecules in the simulation box according to those in a pre-equilibrated system and removes water molecules with atoms that fall within the Van der Waals radii of any atoms of the PAM solutes. Water molecules that were added within a sphere or cylinder of radius 11.25 nm centered at and aligned with the primary axis of the micelle were also removed to avoid unphysical contacts with the hydrophobic core of micelles seeded as spheres or capsules, respectively. This so-called “de-solvated region” disappears well before any measurements are taken and was necessary only for simulation stability during the equilibration process.

Sodium, potassium, and chloride ions were added to the system until a concentration consistent with standard phosphate-buffered saline (PBS) was reached. Additional sodium or chloride ions were then added as necessary to neutralize any net charge. In order to sample states representative of physiological temperature, pH, and pressure, solvated micelles were subject to the following minimization and equilibration procedure: First, the solvated micelle was subject to three steepest-descent minimizations in which (a) all heavy (*i*.*e*., non-hydrogen) atoms were constrained to their starting positions, (b) all amino acid backbone atoms and all carbon atoms within the aliphatic moiety were constrained, and (c) only the carbons most removed from the peptide (the “center atoms”) were constrained.

Having removed any large, nonphysical forces during geometry optimization, the simulated system was brought to thermal equilibrium. This consisted of a multi-step process in which a velocity-rescaling thermostat and Berendsen barostat were applied while the core atoms were fixed in place (Stage 3a). To avoid spurious asymmetries introduced by applying the barostat to a constrained configuration, center of mass reference coordinate scaling was applied (GROMACS mdp file option refcoord-scaling = com). From the final volume of this constrained simulation, an unconstrained equilibration was performed with velocity rescaling every 0.4 ps (Stage 3b). Finally, as the virial, and hence the pressure, was ill-defined in the constrained approach, a subsequent equilibration in which the equations of motion were modified to achieve standard atmospheric pressure and 310 K was performed with the Parrinello–Rahman barostat^61^ and Nosé–Hoover thermostat^62,63^ (Stage 3c).

Simulations were deemed “equilibrated” upon stabilization of both the solvent accessible surface area and either (1) the IUPAP radius of gyration of the entire micelle (given by the trace of the moment of inertia tensor of the micelle) or (2) the radius of gyration about the principal axis for (1) pseudo-spherical or (2) pseudo-cylindrical micelles in accordance with the established approach.^33,64^ Representative data are shown for a PAM comprised of *g*_*k*_ = 70 PAs in Figure 5. After a period of rapid collapse during the constrained tail minimization, the SASA and IUPAP radius of gyration rebound about 10% before settling to the steady-fluctuating values associated with the production run of the simulation (Stage 4a).

**Figure 5:**
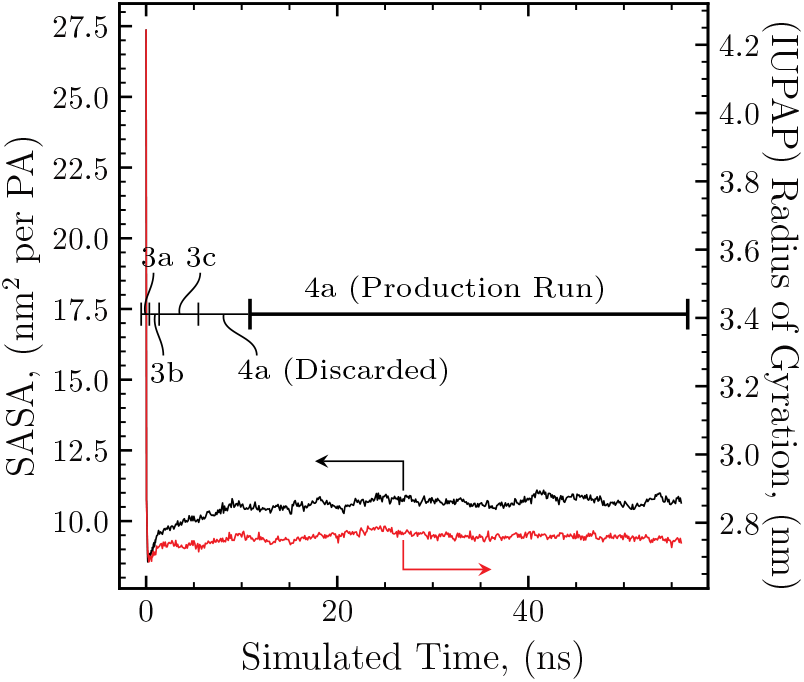
Relaxation of solvent accessible surface area (black line) and radius of gyration (red line) of a simulated PAM of 70 Palm_2_K-NA-(KE)_4_ PAs on the approach to thermal equilibrium. Regions corresponding to the various equilibration stages are indicated. Data from the “production run” portion were used in subsequent analyses.

In this 50 ns production run, the leap-frog algorithm^65^ for integrating Newton’s equations of motion was employed using a 2 fs time step. This was permitted by treating all bonds as constrained via the GROMACS implementation of the LINCS algorithm^66^ (the accuracy of which was controlled via lincs_order = 4 and lincs_iter = 4). Electrostatic interactions were treated with a short-range Coulombic cut off of 1.2 nm, and long-range interactions were handled with the particle-mesh Ewald method with grid spacing of 0.1 nm and fourth-order interpolation (pme_order = 4). To aid in sampling the canonical ensemble, a Nosé–Hoover thermostat was used with a target temperature of 310 K and time constant of 0.4 ps. Periodic boundary conditions were applied to each direction.

### Simulation Analysis

Before analyses were performed, care was taken in placing the micelle in the center of the supercell so that subsequent distance-based analyses could be performed without the aggregate crossing periodic boundaries, and the PAM was confirmed to be intact for the duration of the simulation. The first 5 ns of the “production run” were ascribed to equilibration so the systems could relax to a fixed box size. Configurations were sampled from the remaining production run at 200 ps intervals to approximate independence in consecutive configurations. Such independence was confirmed by decay in the velocity autocorrelation function of the particles comprising the PAM (data not shown). Data were collected from five independently generated simulations of each aggregation number assessed. In particular, 226 configurations were sampled from each production run of micelles seeded with a specified aggregation number and five independently instantiated simulations for each aggregation number were performed. Data across simulations of the same aggregation number were collected for a total of 1130 configurations sampled for each aggregation number. The core of the PAM was assumed to be comprised of both the palmitic acid moieties and the lysine moiety to which they are bonded.

### Mass Weighting

While the total mass of the system does not appear directly in the final expressions for the dimensions of the effective ellipsoid (Equations 5 and 6), the effects of mass weighting persist in the definition of 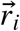, the vector point from the center-of-mass of the PAM to atom *i*. As the goal of this procedure is to characterize the shape of the micelle and not its response to a torque, the procedure presented here with the familiar moment of inertia (MOI) tensor was also modified to assess the influence of mass weighting. Specifically, *m*_*i*_ was taken to be 1 for all particles and 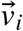 was defined as the vector pointing from the center-of-geometry of the collection of points to the position of the *i*^th^ particle allowing the construction of a pseudo-MOI tensor (pMOI). Taking the reference of the vectors to be the center of geometry rather than the center of mass of the points comprising the micelle differentiates this tensor from the gyration tensor. Equivalence between these tensors is attained for a micelle comprised entirely of particles of equal mass. Whether mass weighting is (MOI) or is not (pMOI) taken into account, this tensor, *I*, provides a unique set of dimensions to define a uniform ellipsoid. Little difference is observed between the characteristic length evaluated from the mass-weighted analysis and non-mass weighted (pMOI) analysis methods (see Figure 6) and as a result, the non-mass weighted analyses were used going forward.

**Figure 6:**
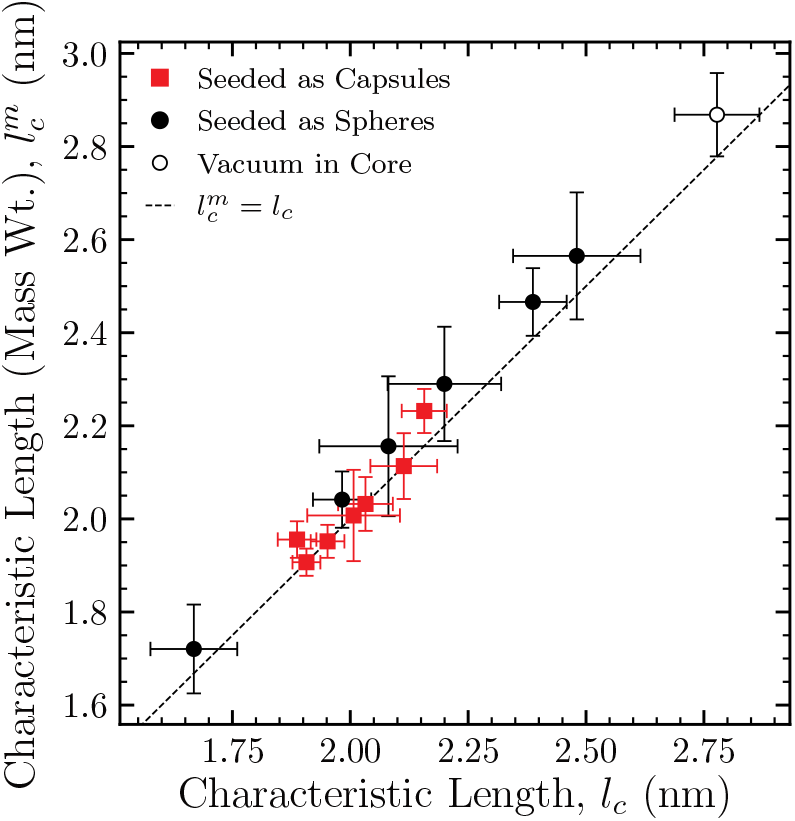
Mass weighting has little influence on the estimation of the characteristic length. A marginal, systematic increase in the mean of the mass-weighted metric for PAMs initially seeded as spheres is perceivable, though the equality condition falls within a single standard deviation, as indicated by the error bars.

In Figure 6, and those to follow, the seeded shape used to generate the micelle in the MD simulation is indicated by the color of the symbol. For the simulations with 99 amphiphiles initially packed into a spherical core, the region of partial vacuum (*i*.*e*., regions of near–zero number density of aliphatic chains) within the micelle core introduced in the equilibration process persisted throughout the simulation (see Figure 7). While the results of the *g*_*k*_ = 99 simulations are likely artifacts, their results are included for two primary reasons. First, this aggregation number appears to coincide with the transition from approximately spherical to more elongated, rod—like micelles. Second, these simulations can be used to infer how trends for primarily spherical micelles would persist beyond the *g*_*k*_ for which they begin to make a transition to more elongated, ellipsoidal morphologies. Interestingly, additional simulations seeded as spheres with *g*_*k*_ > 100 were unstable and the aggregates split apart before any such measures (*e*.*g*., the characteristic length or packing parameter) could be estimated therefrom.

**Figure 7:**
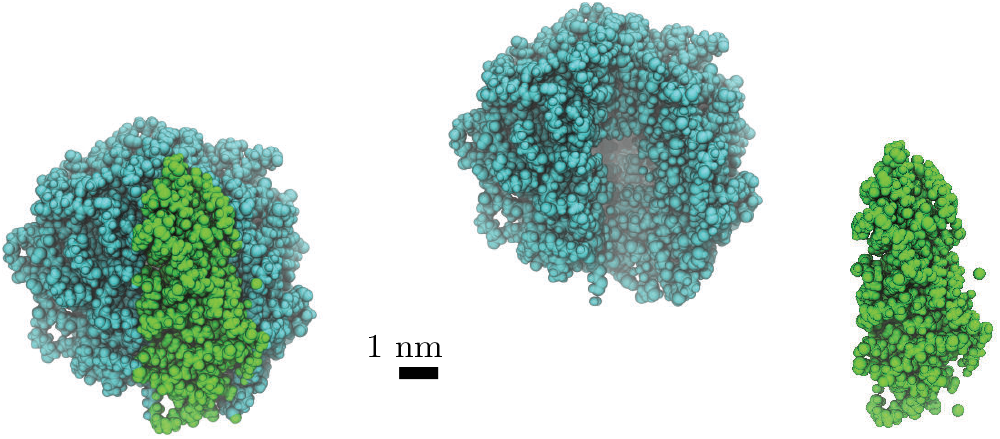
Visualization of non-physical vacuum in the core of the largest simulated spherical micelles.

### Morphology Classification

Morphology classification was done on a “per frame” basis; dimensions describing an ellipsoid (*a, b*, and *c*) were fit to the point mass distribution of species comprising the core for each configuration sampled from the production run. These data were then compiled for each set of simulations of a particular aggregation number. Kernel density estimation was then used to estimate the probability density of an ellipsoid with dimensions (*a, b, c*) in an infinitesimal volume *da db dC* about (*a, b, c*). Specifically, Gaussian kernels were used to form the density estimate on a properly transformed data set via the gaussian_kde function^67^ of the `scipy.stats` module of Python 3.11.4. Applying Silverman’s method for bandwidth selection indicated a value of 0.3 was reasonable for each density constructed in this work.

The (*a, b, c*) values for a given aggregation number were confirmed to be independent of one another when derived from the same simulation and assumed to be so when taken from different simulations of the same aggregation number. Each datum was also assumed to be drawn from the same underlying distribution so that ellipsoid parameters drawn from different frames could be taken to be independent and identically distributed in the estimation of the probability density. Furthermore, when a twenty-one point, fifth-order accurate numerical integration algorithm^68^ subject to these constraints was applied to the density alone, the estimated probability density was found to be properly normalized, thus indicating that all probability density fell within the confines of the convention constraints (*a* ≥ *b* ≥ *c*). A visualization of one such probability density, obtained from simulations of a micelle with *g*_*k*_ = 26, is available in the Supporting Information, wherein, it is evident the parameters are not independent.

Once the probability density for a particular aggregation number, *ρ*_*gk*_, had been found, average shape-dependent properties were estimated for that aggregation number according to the “law of the unconscious statistician” (Equation 33).^69^ For example, for a property *M*,

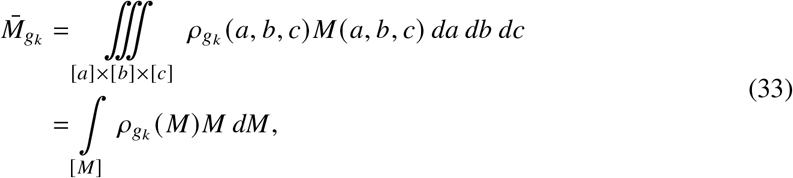

where ∫_[*M*]_ indicates the integral over all relevant *M* values and *ρ*_*gk*_(*M*) is the aggregation-number-dependent probability density of property *M*. In terms of the probability density of the measurable quantities *a, b*, and *c*,

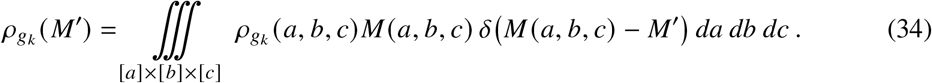

The standard deviation for such properties,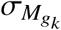, was estimated from the difference in the second raw moment and the square of the first raw moment of the probability density associated with *M*,

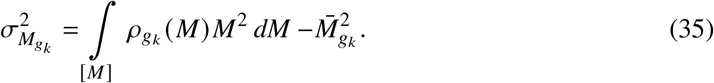

Since we cannot integrate over an infinitely fine mesh, a “pre-limit” form of the Dirac delta function was employed as a function of the parameter *s* defined according to

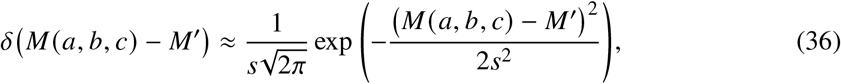

with proper equality restored as *s* → 0. The introduction of this parameter requires a systematic assessment of its choice to ensure its value does not bias the estimate of the probability density. This was accomplished via the selection of a “sufficiently smooth” distribution that still retained its original shape, as described in the Supporting Information.

## Results and Discussion

In this section, results for each measure will be presented in the order they were introduced in the theory section.

### Assessment of Morphology Measures

By analogy to the work of Velinova *et al*.,^45^ the normalized relative shape anisotropy index, 𝒜, is plotted against aggregation number in Figure 8 for simulated micelles of Palm_2_K-NA- (KE)_4_. At low *g*_*k*_, PAM cores are primarily spherical with 𝒜 ≈ 0 and become increasingly anisotropic as *g*_*k*_ increases. A transition region from spherical to rod-like morphologies lies between aggregation numbers of 80 and 160, with further increases in *g*_*k*_ resulting in micelles whose symmetry approaches that of an infinite cylinder.

**Figure 8:**
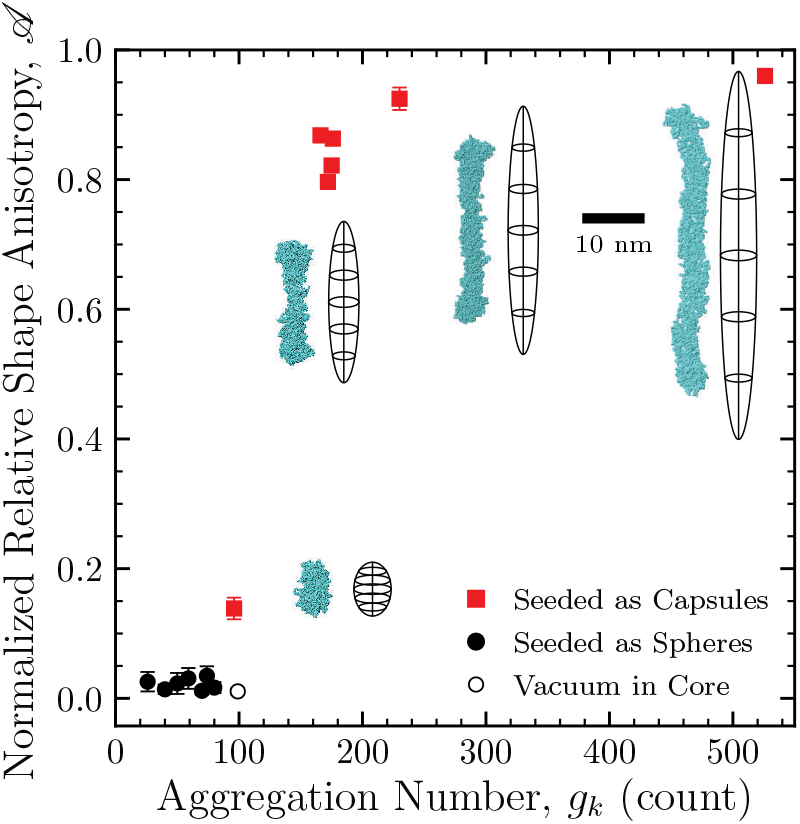
As more amphiphiles are incorporated into a micelle, the normalized relative shape anisotropy of the atoms comprising the core exhibits a transition from spherically symmetric to cylindrically symmetric bodies. Representative visualizations of the PAMs are shown with their best fit ellipsoids and share the indicated scale bar of 10 nm. Error bars indicate a single standard deviation from the mean.

Next, the 𝒜 index and the *P* for the simulated micelles are overlayed on expectations for ideal morphologies in Figure 9. Excellent agreement in the classification of a simulated micelle core as either primarily spherical or primarily cylindrical is observed between 𝒜 and *P*. As expected, both the major and minor aspect ratios of the effective ellipsoid decreased as the morphology changed from spherical to cylindrical. Note that the smallest ratio of the two minor axes of the ellipsoid (*c*/*b* = *α*_2_/*α*_1_ ≈ 0.75) is observed in the midst of the transition region with (*g*_*k*_ = 96). As micelles become larger, this ratio seems to approach a constant value of 0.9. Simulations whose cores were classified as primarily spherical have mean values of *P* ranging from 0.37 to 0.42, whereas those considered to be more rod-like had a *P* range of 0.50 to 0.53. These results indicate that *P* can more sensitively characterize roughly spherical shapes compared to 𝒜, whereas 𝒜 is better able to distinguish more elongated shapes than *P*.

**Figure 9:**
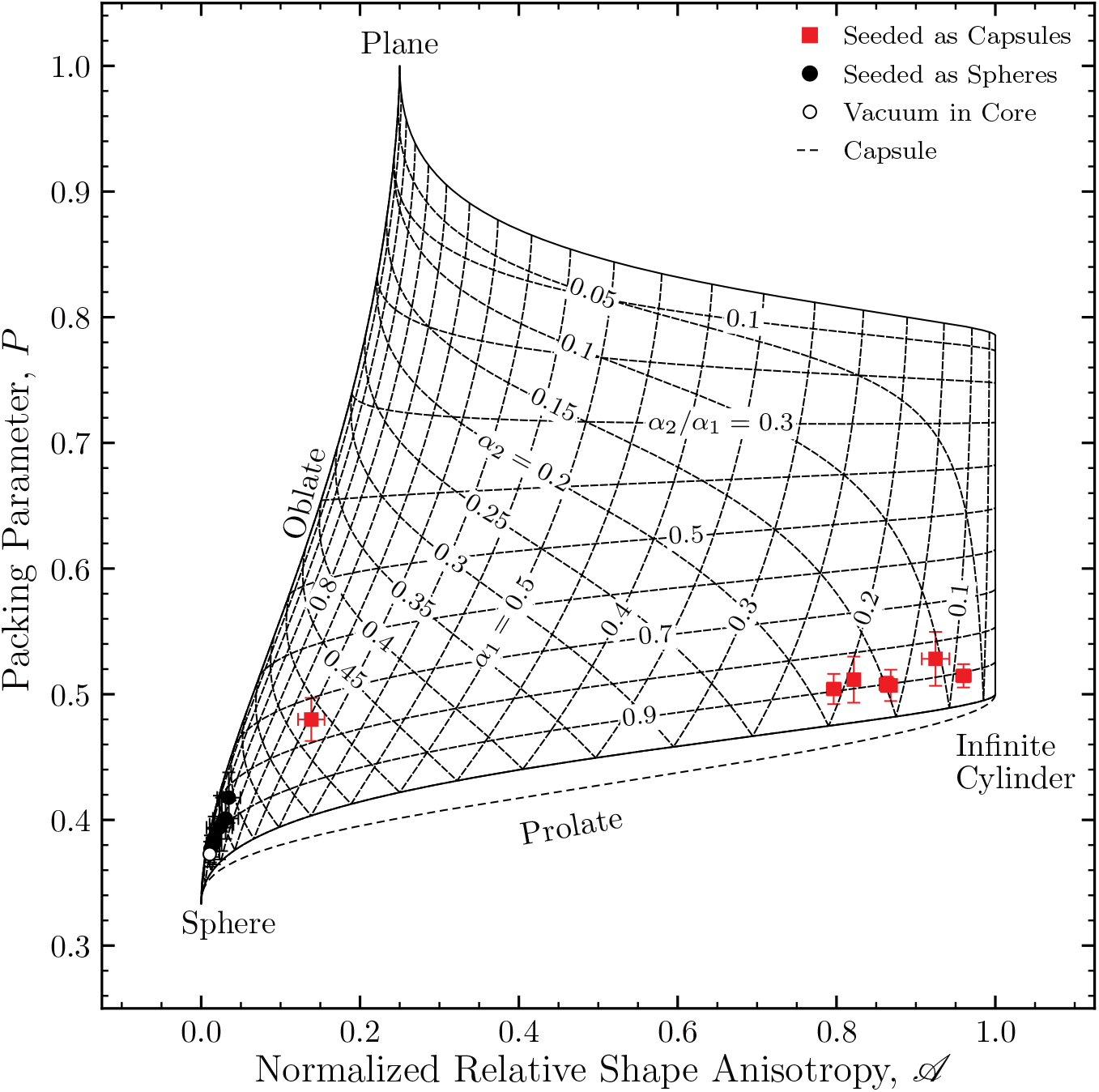
Simulated PAM cores are consistently classified as spheres or cylinders via both the packing parameter and the scaled relative shape anisotropy. As micelles grow (*g*_*k*_ ↑), the ratio between the lengths of the two minor axes of the ellipsoid of best fit decreases for a globular intermediate shape, then approaches a constant, more prolate value for increasingly elongated PAMs. Whereas the packing parameter rapidly approaches 1/2, differences in the scaled relative shape anisotropy are discernable for larger cylindrical micelles.

As the extension of the packing parameter to intermediate morphologies leads to the generalization of the characteristic length, it is imperative to assess this metric against those already established. Figure 10 indicates the values calculated for the characteristic length of Palm_2_KNA-(KE)_4_ PAMs with increasing aggregation number. Previously published results^36^ for MD simulations of hexadecane according to the traditional definition for 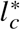 are indicated and are in good agreement with the values calculated for the hydrophobic core of Palm_2_K-NA-(KE)_4_ micelles. The black dashed line in Figure 10 follows from the same arguments used to generate the different *l*_*c*_ values for hexadecane. Assuming the micelle cores formed perfect spheres, *l*_*c*_ = *R*_sph_ and the total volume of a spherical micelle core is given by the sum of the volumes of each hydrophobic portion (*V* = *g*_*k*_ *v*_0_), the expected aggregation number dependence of the *l*_*c*_(that is, the radius for a perfect sphere), is given by

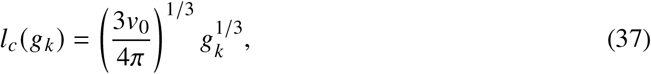

with the volume occupied by the hydrophobic region of an individual PA being the only free parameter. Applying this analysis to the obtained *l*_*c*_ vs. *g*_*k*_ values of the roughly spherical PAM cores (excluding *g*_*k*_ = 99 due to the low density region in the center of the core) results in the indicated estimated volume of 0.38 nm^3^ per chain, corresponding to a molar volume of approximately 230 mL per mole of chains.

**Figure 10:**
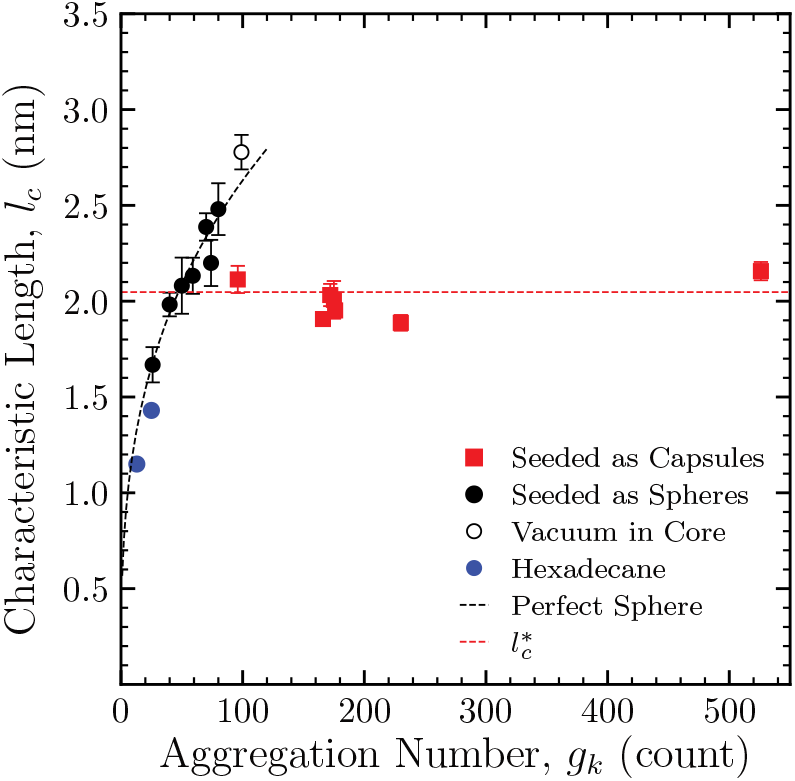
The characteristic length (*l*_*c*_) for Palm_2_K-NA-(KE)_4_ PAMs with varying aggregation number. The *l*_*c*_ for the core of Palm_2_K-NA-(KE)_4_ micelles as defined in Table 2 is in good agreement with theoretical expectations for spherical micelles and empirical results for the maximum extended length. The dashed red line indicates a theoretical maximum extension length for C_16_ alkane chains (*i*.*e*., palmitoyl groups; model described in Ref. 21), and the blue points refer to simulations of spherical hexadecane aggregates (data from Ref. 36). Error bars indicate a single standard deviation from the mean.

The estimated *l*_*c*_ values were also compared to the maximum extended length of hexadecane as given from measurements by Tanford,^21^

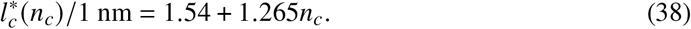

While the *l*_*c*_ values of the PAMs do not strictly abide by this threshold, the peak value attained for spherical micelles just before cylindrical micelles are observed is only 20% larger than this metric. Additionally, there is reason to expect a larger threshold value for *l*_*c*_, as the core of these micelles was defined so as to include the palmitoylated lysine moiety in addition to the alkyl chain. The largest observed *l*_*c*_ corresponds to the metastable spherical micelles that were forced to form with a large aggregation number, and partial vacuum began to appear in the core of the simulated PAMs, suggesting the hydrophobic tails had reached their maximum extension and could not possibly fill the core. Finally, the well-established^23,41,70,71^ trend of the radius of the largest spherical micelles being typically larger than that of the rodlike micelles manifested in our *l*_*c*_ metric.

### Performance of the Packing Free Energy Model

Employing the form of the packing free energy model developed in Equation 32, the free energy change associated with loss of conformational degrees of freedom affiliated with packing the hydrocarbon chains in the core of the micelle was calculated for PAMs simulated with varying aggregation numbers. For PAMs with aggregation numbers below 200 PAs, a distinct aggregation number dependence of the packing free energy is observed (Figure 11). This is consistent with the notion that each subsequent addition of a PA is increasingly penalized as more and more configurational degrees of freedom are lost when more tails are confined to a volume constrained by the length of the fully extended hydrophobic chain. Compared to previously published results for spherical micelles of SDS and DPC,^36^ the values calculated for 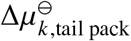 are comparable to those obtained here. Additionally, reasonable quantitative agreement with a published shape-independent model^26^ is observed, providing support for the validity of the shape-dependent model developed here. The shape-independent model posed for the estimation of the packing-free energy depends exclusively on the number of carbons in the aliphatic tail, *n*_*c*_,

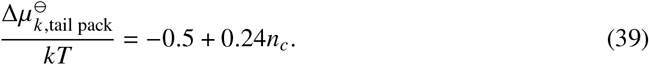

where we have used *n*_*c*_ = 16 and multiplied by a factor of two^20^ to crudely account for the dipalmitoyl moieties chiefly comprising the aliphatic core in these PAMs.

**Figure 11:**
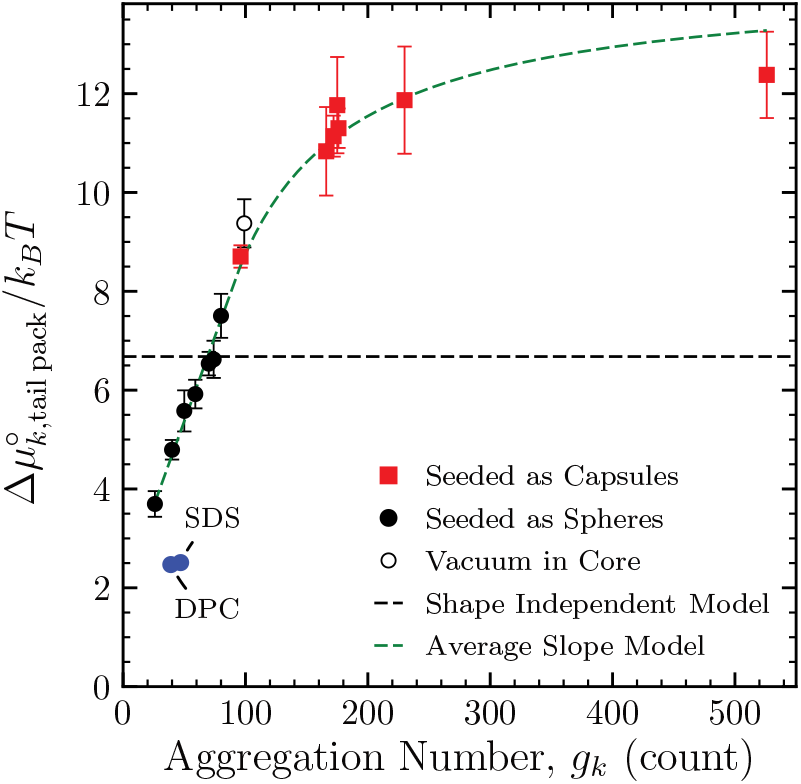
The packing free energy change depends on the shape of the PAM and, for roughly spherical PAMs, the aggregation number. In blue are the results from a published^26^ shape independent model and simulated estimates^36–38^ for the TPFE of two traditional surfactants, sodium dodecyl sulfate (SDS) and dodecylphosphocholine (DPC), which provide context for the feasibility of this model. Error bars indicate a single standard deviation from the mean.

These results are in reasonable agreement with the simple, phenomenological “linear aggregate” model.^23,27,39,41^ In this model, the change in the total tail packing free energy (TPFE) of intermediate, non-regular, finite-sized micelles is estimated by a linear combination of the free energies of micellization of the ideal spherical endcaps and a portion of an infinite cylinder,

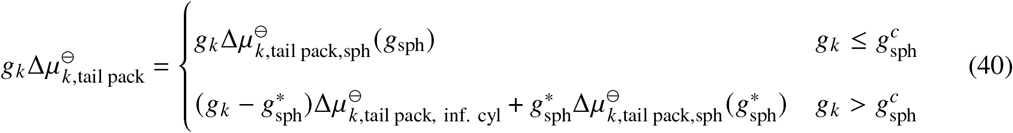

where 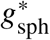 is the aggregation number of the ideal sphere. In rod-like micelles, the free energy penalty associated with packing is expected to saturate as the micelles become increasingly long,

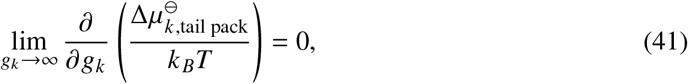

suggesting the total TPFE in this limit is proportional to the aggregation number and 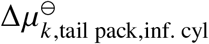, is constant. In roughly spherical micelles, the TPFE per amphiphile is assumed to increase with *g*_*k*_ up to a critical aggregation number 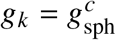 beyond which spherical micelles can no longer form. Lacking the data to resolve each of these parameters, we can make a few simplifying assumptions to reduce the number of parameters. If we assume the critical aggregation number coincides with the ideal sphere, 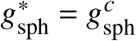 and the aggregation number dependence of the free energy of micellization per amphiphile is a smooth function, the TPFE per amphiphile of an infinite cylinder is fixed,

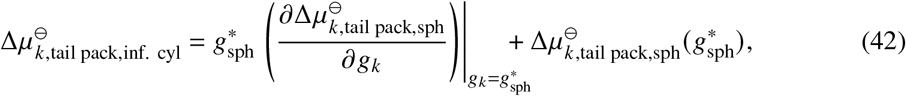

and the TPFE per amphiphile of the simplified linear aggregate model can be approximated with three fitting parameters corresponding to an intercept of the assumed linear spherical region, *b*, the slope of the spherical region, and a critical aggregation number,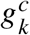. Fitting to the simulation results, the spherical packing free energy increases approximately (0.069 ± 0.005) *k*_*B*_*T* per amphiphile for each additional PA packed into the micelle core. The critical aggregation number coincided nicely with the onset of vacuum within the core of spherical micelle simulations and was estimated to be (90 ± 7) amphiphiles. The intercept parameter was estimated to be (*b* = 1.9 ± 0.3), and the TPFE per amphiphile of the infinite cylinder was (14.3 ± 0.3) *k*_*B*_*T* per amphiphile. This model also captures the increase in the TPFE associated with micelles forced to be in spherical configurations when their *l*_*c*_ is well beyond the optimal value for spherical micelles.

## Conclusions

We have outlined a quantitative means for characterizing the shape of a simulated micelle and successfully applied this approach for MD-simulated PAMs. Specifically, assuming an ellipsoidal shape model whose parameters were defined by the non-mass weighted MOI tensor of the collection of point particles defining the PAM core allowed the precise assignment of shape measures for simulated PAMs whose morphology was in between that of a perfect sphere, an infinite cylinder, or an infinite lamella. These measures are internally consistent as both *P* and 𝒜 clustered to the same morphologies and good visual agreement between the best fit ellipsoid and the visualization of the core was observed.

Because both *P* and 𝒜 are approximately linear at opposite ends of the sphere to infinite cylinder transition (*i*.*e*., it is easier to distinguish spherical bodies from one another than it is to to distinguish cylindrical bodies from each other on the basis of *P*, and vice versa for 𝒜) meaning both measures are useful in distinguishing morphologies. Each of these measures has been expressed in terms of aspect ratios, and we intend to apply these models to estimate the shapes of particles whose aspect ratios can be derived from common characterization techniques, such as transmission electron microscopy (TEM) in future work. Additionally, the extension of the canonical definition of the critical length to the shape-model–dependent *l*_*c*_ results in a metric that is consistent with previous measures and that is applicable to morphologies that lie between those of ideal shapes.

The introduction of a continuous shape model not only allows the quantification of the shape of an arbitrary PAM core but also provides the basis for a tail-packing free energy (TPFE) model. This model demonstrates both quantitative agreement with previous models and adherence to expectations for its limiting behavior.

While the best-fit ellipsoid was presented as the leading choice for the shape model, the implicit convexity of the shape cannot account for the commonly observed^70^ phenomenon of end caps that appear “bulkier than the cylindrical body of the micelle,”^71^ which were also seen in the micelles studied here. Such micelles have core shapes that are near the sphere-to-rod transition. Nonetheless, we wish to emphasize the procedure presented in this work (wherein *𝒜, l*_*c*_, and *P* are defined) can likely be extended to new shape models. For example, for a body with axial symmetry and a desired contour of the micelle core, the surface area of the new shape model can be found from calculus as the surface of revolution produced by the contour, and additional integration(s) can be used to obtain expressions for the volume and MOI for the shape model as outlined further in the Supporting Information.

Because the shape of a nanoparticle plays a crucial role in its function and the current approaches for shape quantification either only apply locally or require the definition of abstract parameters, the shape quantification procedure present in this article provides an important new approach to precisely distinguishing non-ideal shapes. By providing expressions for the generalized packing parameter,*P*, and normalized relative shape anisotropy index, 𝒜, in terms of aspect ratios of an underlying shape model, the shape classification scheme can readily be applied to the analysis of experimental techniques in addition to the analysis of molecular dynamics simulations we demonstrated here. Finally, the flexibility in the formulation of the overall packing parameter facilitates its extension to morphology characterization with a desired shape model.

## Supporting information

Full Supplemental Information

## Acknowledgement

The computation for this work was performed on the high performance computing infrastructure provided by Research Computing Support Services and in part by the National Science Foundation under grant number CNS-1429294 at the University of Missouri, Columbia Missouri.

## Supporting Information Available

Modifications to CHARMM27 force field; full description of statistical methods;

