## Supplementary material for "Generalization of the Packing Parameter for Quantifying the Morphology of Peptide Amphiphile Micelles": Full Supplemental Information

### Modifications to CHARMM27 for Pseudo–Peptide Bonds

In order to make use of GROMACS' built-in topology generator, `pdb2gmx`, the `aminoacids.rtp` file needed to be updated to include the palmitic acid and palmitoylated lysine moieties (e.g., PALM and PALMK). Additionally, GROMACS need to recognize these PALM and PALMK moieties as "protein,"requiring updating of the `residuetypes.dat` file.

Within the `ffbonded.itp` file in the forcefield directory, additional parameters characterizing the pseudo-peptide bonds formed by N-terminal addition of palmitic acid or side chain modified lysine were added to the indicated sections. These parameters were created by copying the analogous term from the original force field and updating the atom types to correspond to those of the new moieties.

```
[ bondtypes ]
; i j   func    b0  kb
CTL2    C   1   0.1538  186188.0 ;
NH1 CL  1   0.1344  309616.0 ;
[ angletypes ]
; i j   k   func    th0 cth ub0 cub
CT1 NH1 CL  5   120.0000    418.4   0.0 0.0
CT2 NH1 CL  5   120.0000    418.4   0.0 0.0
H   NH1 CL  5   123.0000    284.512 0.0 0.0
HA  CTL2    C   5   109.50  276.144 0.2163  25104.0
NH1 C   CTL2    5   116.5000    669.44  0.0 0.0
NH1 CL  CTL2    5   116.5000    669.44  0.0 0.0
O   C   CTL2    5   121.0000    669.44  0.0 0.0
OCL CL  NH1 5   122.5000    669.44  0.0 0.0
HAL2    CTL2    C   5   110.10  221.752 0.2179  18853.104
C   CTL2    CTL2    5   113.50  488.2728    0.2561  9338.688
[ dihedraltypes ]
CTL2    CL  NH1 CT2 9   180.00  10.46   2
HA  CT2 NH1 CL  9   0.00    0.0 3
OCL CL  NH1 CT1 9   180.00  10.46
OCL CL  NH1 CT2 9   180.00  10.46
OCL CL  NH1 H   9   180.00  10.46
OCL NH1 CL  CTL2    2   0.0000  1004.16
OCL NH1 CTL2    CL  2   0.0000  1004.16
```

Listing S1: Edits to ffbonded.itp

### Probability Density Estimation and Rendering

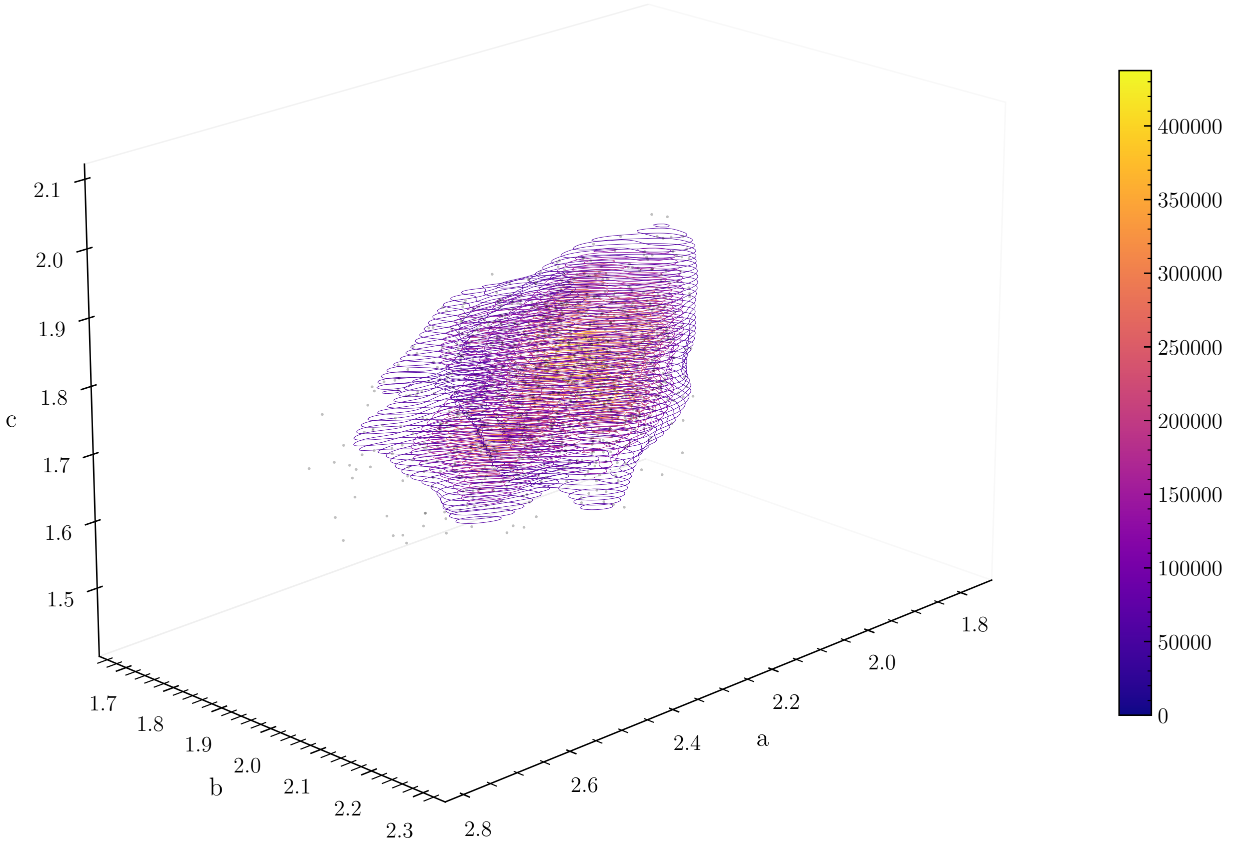


Figure S1: 3D rendering of the probability density, $\rho_{g_k}(a, b, c)$ estimated via kernel density estimation for the core of simulated PAMs with aggregation number, $g_k = 26$. Grid spacing is $\Delta a = 6.611 \times 10^{-3}$ nm, $\Delta b = 4.195 \times 10^{-3}$ nm, and $\Delta c = 6.958 \times 10^{-3}$ nm. Contours are plotted at constant $c$, and gray points indicate the individual values of $(a, b, c)$ calculated from a specific frame of a simulation.

#### Distribution Selection

The probability densities of specific properties of interest, $M$, from the kernel density derived probability densities of "measurable" parameters $a$, $b$, and $c$ are given analytically by Equation (43) in the main text,

$$\rho_{g_k}(M') = \iiint\limits_{[a]\times[b]\times[c]} \rho_{g_k}(a,b,c)M(a,b,c)\delta\left(M(a,b,c) - M'\right) da\, db\, dc\,. \tag{43}$$

Functionally, since we cannot integrate over an infinitely fine mesh, the "pre-limit" form of the Dirac delta function as a function of the parameter $s$ defined according to Equation (45) in the main text,

$$\delta(M(a,b,c) - M') \approx \frac{1}{s\sqrt{2\pi}} \exp\left(-\frac{(M(a,b,c) - M')^2}{2s^2}\right), \tag{45}$$

is used to aid in the evaluation of this integral.

$$\rho_{g_k}(M') = \lim_{s\to 0} \frac{1}{s\sqrt{2\pi}} \iiint\limits_{[a]\times[b]\times[c]} \rho_{g_k}(a,b,c)M(a,b,c)e^{\left(-\frac{(M(a,b,c)-M')^2}{2s^2}\right)} da\, db\, dc \tag{S1}$$

Then, the triple integral is approximated with a fifth order accurate numerical integration algorithm as follows,[S1]

1. The domain of each variable, $x \subset (a,b,c)$, was assumed to be $\bar{x} \pm 5 \times s_x$ where $\bar{x}$ and $s_x$ are the mean and sample standard deviations calculated directly from the set of variables obtained from each frame.

2. Each domain was then split into $N$ equal parts, forming $N^3$ "cubelets" of side lengths $\Delta a = \frac{10}{N}s_a$, $\Delta b = \frac{10}{N}s_b$, and $\Delta c = \frac{10}{N}s_c$.

3. The integrand function was then evaluated at twenty-one non-equally weighted points within each cubelet as indicated in the original source.[S1]

4. The value of the integral over a cubelet whose origin was located at $(N_a, N_b, N_c)$ could then

be estimated as,

$$
\begin{aligned}
I(N_a, N_b, N_c) &= \int_0^{\Delta c} \int_0^{\Delta b} \int_0^{\Delta a} f(a', b', c')\, da'\, db'\, dc' \\
&\approx \frac{\Delta a \Delta b \Delta c}{8 \times 45} \Bigg\{ -496 f\left(\Delta a \left(N_a + \tfrac{1}{2}\right), \Delta b \left(N_b + \tfrac{1}{2}\right), \Delta c \left(N_c + \tfrac{1}{2}\right)\right) \\
&+ 128 \Bigg[ f\left(\Delta a \left(N_a + \tfrac{1}{2}\right), \Delta b \left(N_b + \tfrac{1}{2}\right), \Delta c \left(N_c + \tfrac{1}{4}\right)\right) \\
&\quad + f\left(\Delta a \left(N_a + \tfrac{1}{2}\right), \Delta b \left(N_b + \tfrac{1}{4}\right), \Delta c \left(N_c + \tfrac{1}{2}\right)\right) \\
&\quad + f\left(\Delta a \left(N_a + \tfrac{1}{4}\right), \Delta b \left(N_b + \tfrac{1}{2}\right), \Delta c \left(N_c + \tfrac{1}{2}\right)\right) \\
&\quad + f\left(\Delta a \left(N_a + \tfrac{1}{2}\right), \Delta b \left(N_b + \tfrac{1}{2}\right), \Delta c \left(N_c + \tfrac{3}{4}\right)\right) \\
&\quad + f\left(\Delta a \left(N_a + \tfrac{1}{2}\right), \Delta b \left(N_b + \tfrac{3}{4}\right), \Delta c \left(N_c + \tfrac{1}{2}\right)\right) \\
&\quad + f\left(\Delta a \left(N_a + \tfrac{3}{4}\right), \Delta b \left(N_b + \tfrac{1}{2}\right), \Delta c \left(N_c + \tfrac{1}{2}\right)\right) \Bigg] \\
&+ 8 \Bigg[ f\left(\Delta a \left(N_a + \tfrac{1}{2}\right), \Delta b \left(N_b + \tfrac{1}{2}\right), \Delta c N_c\right) \\
&\quad + f\left(\Delta a \left(N_a + \tfrac{1}{2}\right), \Delta b N_b, \Delta c \left(N_c + \tfrac{1}{2}\right)\right) \\
&\quad + f\left(\Delta a N_a, \Delta b \left(N_b + \tfrac{1}{2}\right), \Delta c \left(N_c + \tfrac{1}{2}\right)\right) \\
&\quad + f\left(\Delta a \left(N_a + \tfrac{1}{2}\right), \Delta b \left(N_b + \tfrac{1}{2}\right), \Delta c \left(N_c + 1\right)\right) \\
&\quad + f\left(\Delta a \left(N_a + \tfrac{1}{2}\right), \Delta b \left(N_b + 1\right), \Delta c \left(N_c + \tfrac{1}{2}\right)\right) \\
&\quad + f\left(\Delta a \left(N_a + 1\right), \Delta b \left(N_b + \tfrac{1}{2}\right), \Delta c \left(N_c + \tfrac{1}{2}\right)\right) \Bigg] \\
&+ 5 \Big[ f\left(\Delta a \left(N_a + 1\right), \Delta b N_b, \Delta c N_c\right) + f\left(\Delta a N_a, \Delta b (N_b + 1), \Delta c N_c\right) \\
&\quad + f\left(\Delta a N_a, \Delta b N_b, \Delta c (N_c + 1)\right) + f\left(\Delta a N_a, \Delta b N_b, \Delta c N_c\right) \\
&\quad + f\left(\Delta a (N_a + 1), \Delta b (N_b + 1), \Delta c N_c\right) + f\left(\Delta a N_a, \Delta b (N_b + 1), \Delta c (N_c + 1)\right) \\
&\quad + f\left(\Delta a (N_a + 1), \Delta b N_b, \Delta c (N_c + 1)\right) \\
&\quad + f\left(\Delta a (N_a + 1), \Delta b (N_b + 1), \Delta c (N_c + 1)\right) \Big] \Bigg\}
\end{aligned}
\tag{S2}
$$

Finally, the density is estimated as a function of the parameter $s$ and the number of cubelets, $N$ by,

$$\rho_{g_k}(M'; s, N) \approx \frac{1}{s\sqrt{2\pi}} \sum_{N_a=0}^{N-1} \sum_{N_b=0}^{N-1} \sum_{N_c=0}^{N-1} I(N_a, N_b, N_c; s). \tag{S3}$$

and is subject to the normalization condition,

$$\int_{\bar{M}-5s_M}^{\bar{M}+5s_M} \rho_{g_k}(M'; s, N) dM' = 1 \tag{S4}$$

Evidently, this estimate of the probability density depends on both the parameter $s$ and $N$. Choice of these parameters was made via the selection of a "sufficiently smooth" distribution that still retained it original shape and the "smoothness" of the distribution was quantified via a scaled, discrete Lipschitz constant, called the the relative roughness, $RR$,

$$RR = \frac{\sup \left| \frac{\Delta\rho(M)}{\Delta M} \right|}{\frac{\rho(M^*)}{M^*}} \tag{S5}$$

where $M^*$ is the value of the property of interest (e.g., the characteristic length or the packing parameter) where the probability density, $\rho$ is the highest: the mode of the distribution. The shape of a given density was characterized on the basis of its first few moments, the interquartile range, and combinations thereof. Namely, the skew, excess kurtosis, bimodality coefficient, and the normalized inter-quartile range (IQR) were plotted against the $RR$ parameter and a metric $Q = 1/sN$ to assess for convergence of the integration. To aid in the presentation of the high dimensional data, values of the parameters $s$ and $N$ were indicated by their color according to Figure S2. Figure S3 indicates the dependence of probability density on its parameters as evaluated by the aforementioned metrics. As expected, $RR$ decreases with increasing $s$, though complete characterization of the shape of the distribution requires the specification of both $s$ and $N$. As $Q$ decreases at a constant $s$ value, the accuracy of the integration is improved at first, but eventually is oversmoothed. Fortunately, however, the balance between integration accuracy and smoothness of the distribution need not be determined to the nth degree as convergence of both the mean and standard deviations of the probability density is attained for multiple sets of these parameters (see Figure S4). The convergence of the probability density itself is shown in Figure S5.

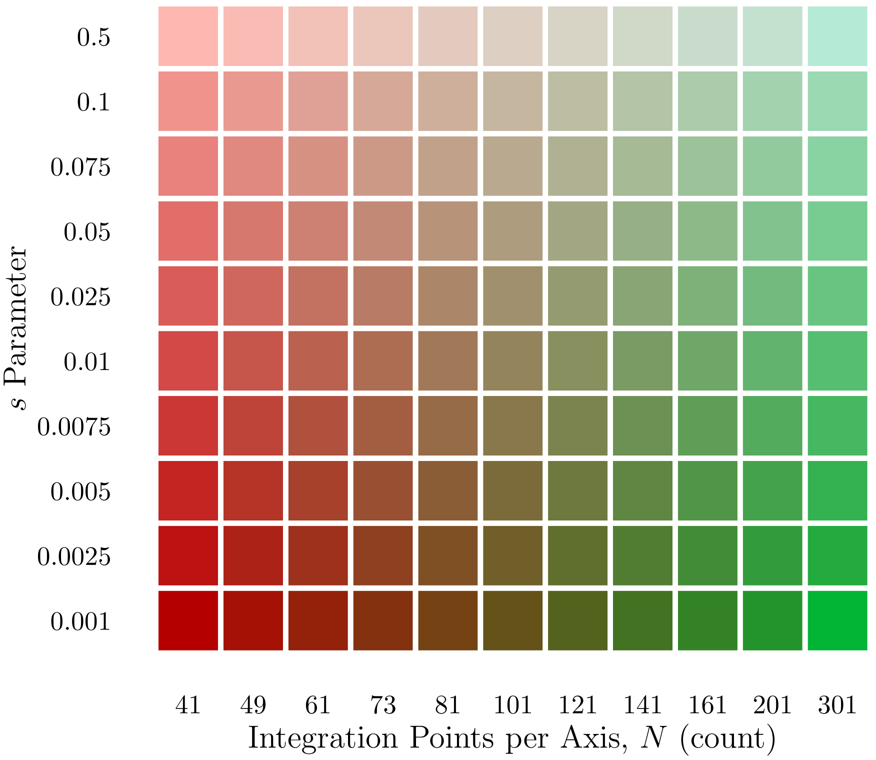


Figure S2: Color key use to indicate the parameters used to estimate a probability density. Colors soften from red or green to pastel variants as the gaussian approximation to the Dirac delta distribution is softened. Red transitions through brown into green colors as the integration resolution increases.

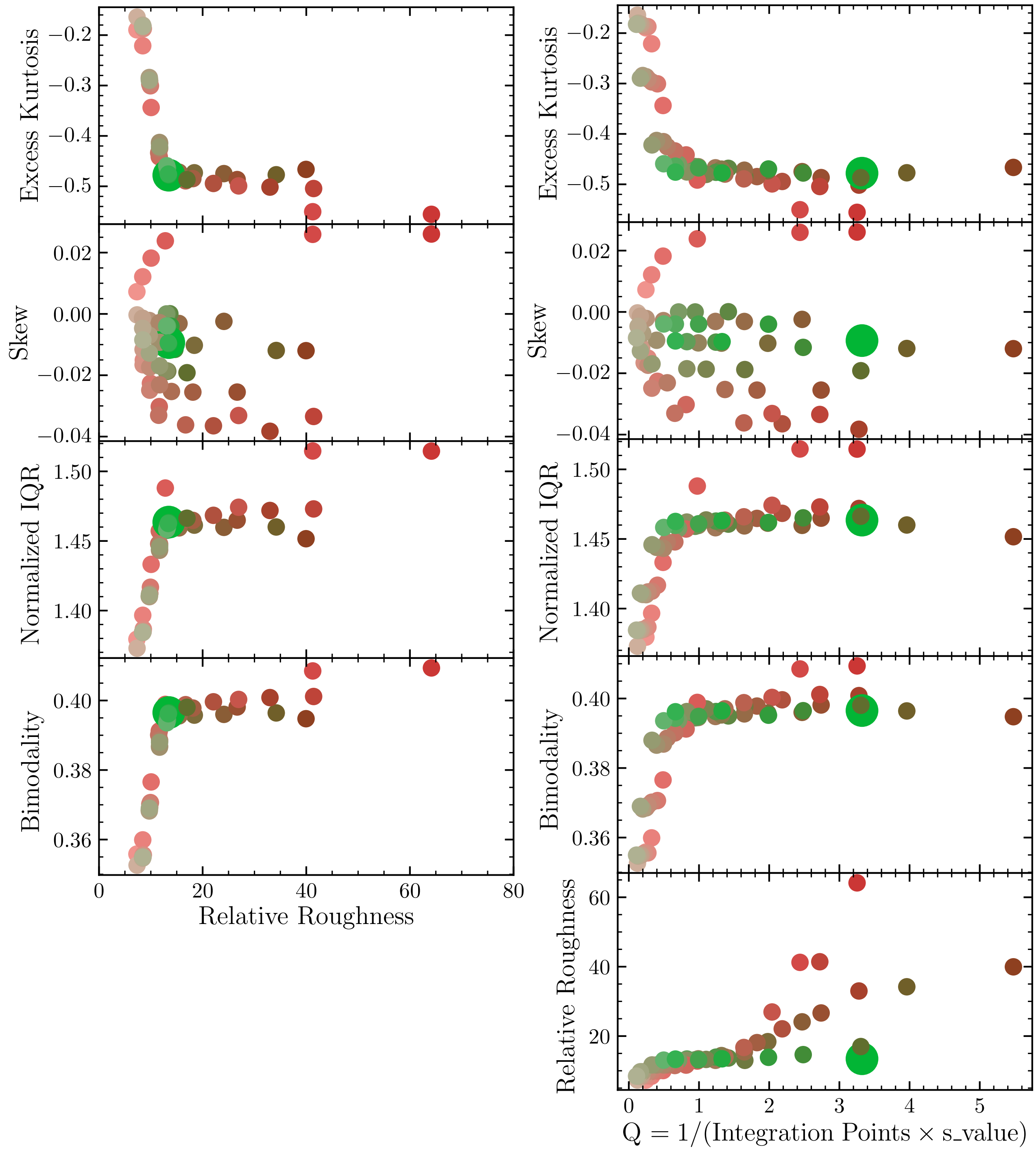


Figure S3: Probability density shape and smoothness is dependent on coupled $s$ and $N$ parameters. The larger point indicates the parameter set used to estimate the mean and standard deviation of $l_c$ of a micelle with $g_k = 26$.

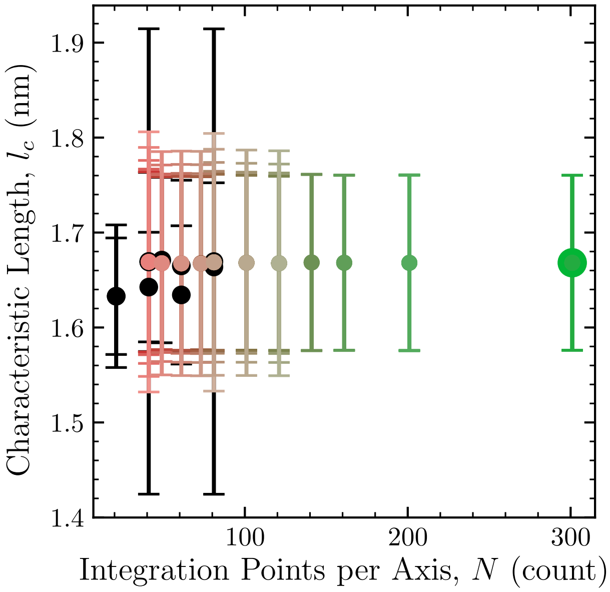


Figure S4: Convergence in the estimation of the mean characteristic length is dependent on the value of $s$ and $N$. Parameters are indicated by the color of the point according to Figure S2 and black points have $RR > 80$. Error bars indicated the standard deviation of the estimated probability density. The parameter set for the larger point was used to estimate the $l_c$ distribution, though convergence in the mean and standard deviation was achieved in parameter sets with larger $s$ and smaller N.

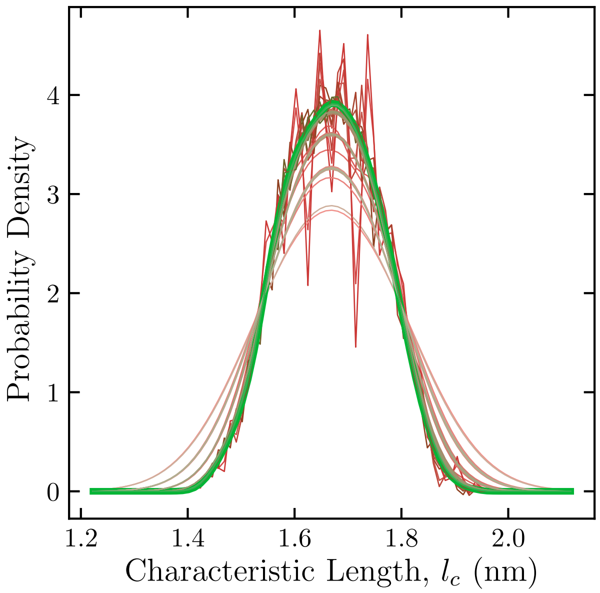


Figure S5: Probability density of the characteristic length for a PAM with $g_k = 26$. The probability density from which the mean and standard deviation of the characteristic length of a micelle with $g_k = 26$ is thicker than the other density estimates.

### Example Extension to A Solid of Revolution

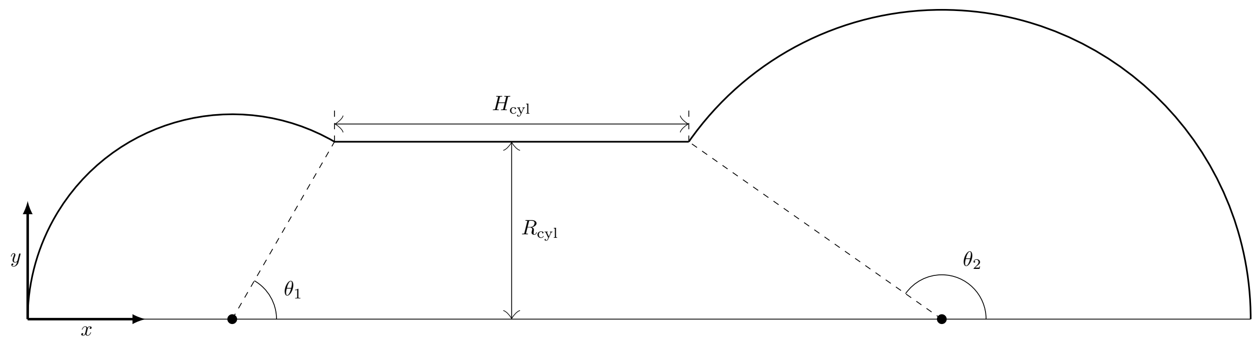


Figure S6: Example plane shape for an axially symmetric body

Figure S6 indicates an example "plane shape" (i.e., 2-D shape) that, upon revolution about the $x$-axis might be used to model a system. In particular, this example indicates a cylindrical region capped by two semi-spheres of different radii: $R_{\mathrm{cyl}}/\sin\theta_1$ and $R_{\mathrm{cyl}}/\sin\theta_2$, respectively. The expression for the function characterizing the edge of the shape (or surface upon revolution) can be expressed as a piece-wise function of the indicated parameters: $\theta_1$, $\theta_2$, $R_{\mathrm{cyl}}$, and $H_{\mathrm{cyl}}$,

$$f(x;\theta_1,\theta_2,R_{\mathrm{cyl}},H_{\mathrm{cyl}}) = \begin{cases} \sqrt{\left(\frac{R_{\mathrm{cyl}}}{\sin\theta_1}\right)^2 - \left(x - \frac{R_{\mathrm{cyl}}}{\sin\theta_1}\right)^2} & 0 \le x \le x_1 \\ R_{\mathrm{cyl}} & x_1 < x < x_2 \\ \sqrt{\left(\frac{R_{\mathrm{cyl}}}{\sin\theta_2}\right)^2 - (x - x_4)^2} & x_2 < x \le x_3 \end{cases} \tag{S6}$$

where $x_1 = \frac{R_{\mathrm{cyl}}}{\sin\theta_1}(1+\cos\theta_1)$, $x_2 = x_1 + H_{\mathrm{cyl}}$, $x_3 = x_2 + \frac{R_{\mathrm{cyl}}}{\sin\theta_2}(1-\cos\theta_2)$, and $x_4 = x_3 + \cos\theta_2$. From this function, the surface area, $A$, and volume, $V$, of the revolved body are given from calculus,

$$A(\theta_1,\theta_2,R_{\mathrm{cyl}},H_{\mathrm{cyl}}) = \int_0^{x_3} f(x;\theta_1,\theta_2,R_{\mathrm{cyl}},H_{\mathrm{cyl}})\sqrt{1+\left(\frac{df}{dx}\right)^2}\,dx, \tag{S7}$$

and

$$V(\theta_1,\theta_2,R_{\mathrm{cyl}},H_{\mathrm{cyl}}) = \int_0^{x_3} f(x;\theta_1,\theta_2,R_{\mathrm{cyl}},H_{\mathrm{cyl}})^2\,dx\,. \tag{S8}$$

Assuming a uniform density throughout the, the moment of inertia about the $x$-axis is given by

$$I_x(\theta_1, \theta_2, R_{\text{cyl}}, H_{\text{cyl}}) = \frac{\pi}{2}\frac{M}{V}\int_0^{x_3} f(x; \theta_1, \theta_2, R_{\text{cyl}}, H_{\text{cyl}})^4 \, dx, \tag{S9}$$

and from the parallel-axis theorem, the moment of inertia about the $y$ and $z$-axes are given by

$$I_y(\theta_1, \theta_2, R_{\text{cyl}}, H_{\text{cyl}}) = \frac{I_x}{2} + \frac{\pi}{2}\frac{M}{V}\int_0^{x_3} x^2 f(x; \theta_1, \theta_2, R_{\text{cyl}}, H_{\text{cyl}})^2 \, dx \,. \tag{S10}$$

While the moment of inertia will give a clear value for $\mathscr{A}$, it is clearly not a unique value. That is, multiple sets of the parameters will have the same value of the normalized relative shape anisotropy index. In a similar fashion, the four parameters in this shape model cannot be uniquely defined from the eigenvalues of the MOI tensor.

However, progress can be made by assuming the value of a particular parameter, or on a unitless basis, the ratio of two model parameters (e.g., $H_{\text{cyl}}/R_{\text{cyl}}$. Then, as in the formulation of the main text, establishing the boundary conditions (i.e., limiting behavior) would be required to find expressions for the characteristic length and packing parameter. In this example, one such boundary condition would require the characteristic length of the solid of revolution to be equal to that of the capsule for all positive values of $R_{\text{cyl}}$ and $H_{\text{cyl}}$ when $\theta_1 = \theta_2 = \pi/2$.

While estimation of the index $\mathscr{A}$ should be possible from this procedure (assuming the necessary integrations can be performed), arriving at unique expressions for $P$ and $l_c$ that apply to the entire domain of the parameter space is contingent upon the ability to define a sufficient number of independent boundary conditions ($\geq$ number of parameters $+1$).
